# The microprotein Minion controls cell fusion and muscle formation

**DOI:** 10.1101/122697

**Authors:** Qiao Zhang, Ajay Vashisht, Jason O’Rourke, Stéphane Y. Corbel, Rita Moran, Angelica Romero, Loren Miraglia, Jia Zhang, Eric Durrant, Christian Schmedt, Srinath C. Sampath, Srihari C. Sampath

## Abstract

Although recent evidence has pointed to the existence of small open reading frame (smORF)-encoded microproteins in mammals, the functional repertoire of this microproteome remains to be determined^1^. In skeletal muscle, proper development requires fusion of mononuclear progenitors to form multinucleated myotubes, a critical but poorly understood process^2,3^. Here we report the identification of a small ORF encoding an essential skeletal muscle specific microprotein we term Minion (*m*icroprotein *in*ducer of fusion). Myogenic progenitors lacking Minion differentiate normally but fail to form syncytial myotubes, and Minion-deficient mice die perinatally with marked reduction in fused muscle fibers. This fusogenic activity is conserved to the human Minion ortholog, previously annotated as a long noncoding RNA. Loss-of-function studies demonstrate that Minion is the factor providing muscle specific fusogenic function for the transmembrane protein Myomaker^4^. Remarkably, we demonstrate that co-expression of Minion and Myomaker is sufficient to induce rapid cytoskeletal rearrangement and homogeneous cellular fusion, even in non-muscle cells. These findings establish Minion as a novel microprotein required for muscle development, and define a two-component program for the induction of mammalian cell fusion, enabling both research and translational applications. Importantly, these data also significantly expand the known functions of smORF-encoded microproteins, an under-explored source of proteomic diversity.

---

In addition to canonically defined protein-coding genes, recent studies have indicated the existence of a new class of mammalian genes^5^. These small open reading frames (smORFs) are transcribed and translated by usual means, but are largely unrecognized as protein-coding genes by virtue of their size, typically encoding microproteins <100 amino acids (aa) in length^1^. Although estimates vary widely, the human and mouse genomes are thought to contain at least several thousand of these “hidden” protein-coding genes^1^. Intriguingly, of the small number of currently known mammalian microproteins, several have been identified in muscle^6-10^. These largely encode regulatory factors for the sarco/endoplasmic reticulum Ca^2+^-ATPase (SERCA), with structural similarity to known SERCA-regulatory proteins such as sarcolipin and phospholamban^6-8^. Of note however, no essential mammalian microprotein has been described.

Muscle development requires temporally regulated stem cell activation and differentiation, fusion of progenitors to form syncytial myotubes, and maturation of myotubes to generate contractile myofibers. While the early and late stages of this process have been intensively studied^2,11^, our understanding of the mechanisms and regulatory factors controlling cell fusion remains incomplete, particularly in mammals^3,12^. A recent major advance was the identification of the transmembrane protein Tmem8c/myomaker, which is necessary for myoblast fusion and sufficient for fusion of non-muscle cells to differentiating muscle. Importantly however, myomaker expression alone cannot induce fusion of non-muscle cells with one another, suggesting the existence of one or more additional factors that are expressed in differentiating muscle cells and required to drive *de novo* cell fusion^4,13^.

To identify novel microproteins playing key roles in skeletal muscle, we performed whole transcriptome RNA-seq analysis of uninjured and regenerating muscle. We specifically sought to identify transcripts demonstrating strong temporal regulation, annotated ORF length of less than 100 codons, and a corresponding dynamic pattern of transcriptional regulation during mouse myoblast differentiation *in vitro* (Fig. 1a, left). We focused on gene regulation at day 3 post injury in order to exclude effects related to the immediate post-injury immune response. The predicted gene 7325 (*gm7325*) was the only gene meeting all criteria, encoding a putative 84 aa microprotein with possible expression in ES cells but no known function^14^. For reasons described below, we named this gene *minion* (*m*icroprotein *in*ducer of fus*ion*). The temporal pattern of *minion* expression was distinct from that of two other smORFs, but notably was similar to that of myomaker (Fig. 1a, right)^4^.

**Fig. 1.**
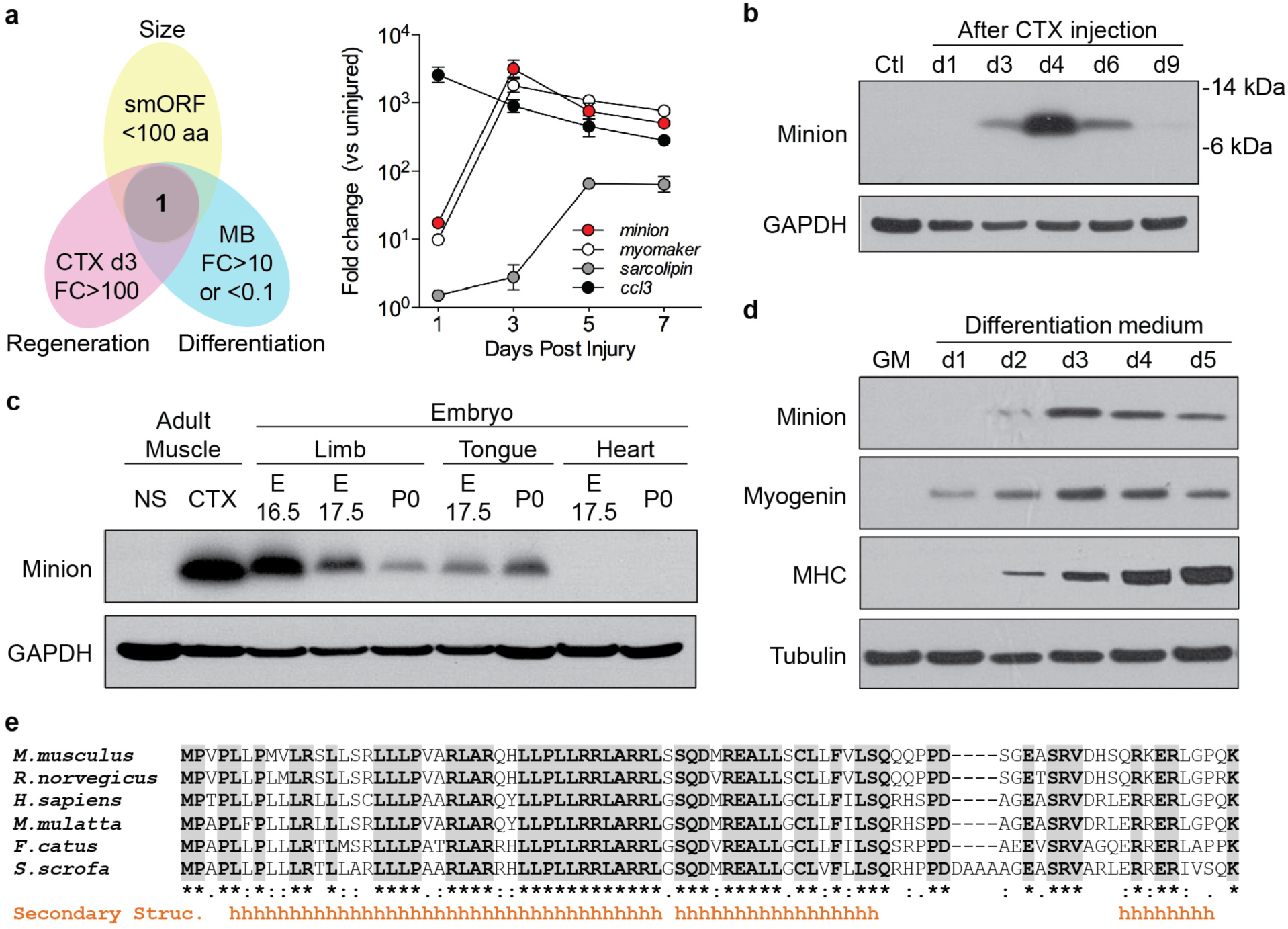
The microprotein minion is skeletal muscle specific and highly expressed during development and regeneration. **(a)** Left: Overlap of RNA-seq from regenerating adult mouse *tibialis anterior* (TA) muscle and differentiating C2C12 myoblasts (MB). CTX, cardiotoxin; FC: fold change compared to uninjured muscle (bottom left) or undifferentiated myoblasts (bottom right). Right: fold change of reads per kilobase per million mapped reads (RPKM) for selected genes upregulated after CTX injury. Values are normalized to uninjured muscle. Mean ± SD of fold change, n=3 per time point. **(b)** Western blot of control uninjured (Ctl) and CTX-injured regenerating adult TA muscle (n=4). **(c)** Western blot of embryonic muscle samples (n=3). Day 4 post-CTX TA or normal saline (NS) injection were positive and negative controls. E, embryonic day; P: post-natal. **(d)** Western blot of C2C12 myoblasts cultured under growth conditions (GM) or under differentiation conditions (DM) for the indicated number of days (n=4). MHC, myosin heavy chain. (b-d) Glyceraldehyde-3-phosphate dehydrogenase (GAPDH) and Tubulin served as loading controls. **(e)** Protein sequence alignment of mouse minion with putative orthologs from other mammalian species (GenBank accession numbers and UniProt IDs from top to bottom are: NP_001170939.1, XP_017452417.1, NP_001302423.1, EHH18375.1, M3X8W7_FELCA, and F1RQU5_PIG.)

Western blot confirmed that the *minion* transcript is translated; minion protein was absent in uninjured tibialis anterior (TA) muscle but strongly induced during regeneration, peaking three to four days following injury (Fig. 1b). Immunofluorescence analysis demonstrated minion expression within nascent regenerating myofibers (Extended Data Fig. 1a), whereas minion protein was not detectable in uninjured adult muscle (Extended Data Fig. 1b) nor in multiple additional non-muscle tissues (Extended Data Fig. 1c). RNA-seq analysis of early embryonic development revealed *minion* expression which was detectable as early as somite stage 15 but greatly increased by somite stage 36, following limb and tail bud formation (Extended Data Fig. 1d). Expression of *minion* was seen in embryonic skeletal muscle of both somitic (limb, tongue) and non-somitic (extraocular and facial muscles) origin, but importantly not in embryonic or neonatal heart muscle (Fig. 1c, Extended Data Fig. 1e). Both mRNA and protein levels of minion increased rapidly during *in vitro* myoblast differentiation (Fig. 1d and Extended Data Fig. 2a-c). Although the full-length minion protein is predicted to contain an N-terminal signal sequence and predominant alpha-helical secondary structure (Fig. 1e), overexpression and supernatant concentration demonstrated no evidence of protein secretion (Extended Data Fig. 2d). Subcellular fractionation did however confirm significant enrichment within the membrane-associated fraction containing plasma membrane, ER, and Golgi, suggesting insertion into or association with a membrane compartment (Extended Data Fig. 2e).

TBLASTN search revealed a putative human *Minion* homolog (*hMinion*) with an intact ORF of 84 codons (Extended Data Fig. 3a), despite prior annotation of the transcript as a long noncoding RNA (RP1-302G2.5; LOC101929726). Evolutionary conservation of the protein coding sequence was seen across mammalian species (Fig. 1e), however no convincing sequence homolog was found in *Drosophila* or other invertebrate species. No amino acid sequence similarity was seen to sarcolipin, phospholamban, or the recently reported microprotein DWORF ^6^. We noted that minion expression during muscle cell differentiation slightly trailed that of the basic helix-loop-helix transcription factor Myogenin (Fig. 1d), suggesting control by canonical muscle regulatory factors (MRFs, e.g. MyoD and Myogenin). Indeed, analysis of the upstream regulatory regions of human and mouse *minion* loci revealed evolutionarily conserved E-box binding sites for MRFs (Extended Data Fig. 3a). Both MyoD and Myogenin specifically bound these sites in differentiating myoblasts, as shown by ENCODE whole genome ChIP-seq (Extended Data Fig. 3b)^15^. We conclude that minion is an uncharacterized but conserved, membrane-associated, and skeletal muscle-specific microprotein.

The spatial and temporal pattern of minion expression together with the presence of functional MRF-binding E-boxes strongly suggested a role for minion in skeletal muscle development. To test this, we used CRISPR/Cas9 genome editing to generate *minion*-deficient mice (Fig. 2a). Two guide RNAs (gRNAs) targeting the single coding exon were coinjected into embryos, and F_0_ pups were screened for mutations. Small insertion/deletion mutations as well as larger deletions between gRNA target sites were identified (Extended Data Fig. 4a), and a founder line containing a 135 bp in-frame deletion (*minion*^Δ/Δ^; Fig. 2a, b, Extended Data Fig. 4b) was characterized further. Although heterozygous *minion*^Δ^*^/^*^+^ animals were viable and recovered at expected Mendelian ratios, live homozygous mutant *minion*^Δ/Δ^ animals could only be observed prenatally, and no viable neonatal or adult *minion*^Δ/Δ^ animals were recovered (Extended Data Fig. 4c, d). Loss of minion protein was confirmed in both embryonic and perinatal limb and tongue skeletal muscle from *minion*-deficient animals (Extended Data Fig. 5). These findings are consistent with perinatal lethality in the absence of minion.

**Fig. 2.**
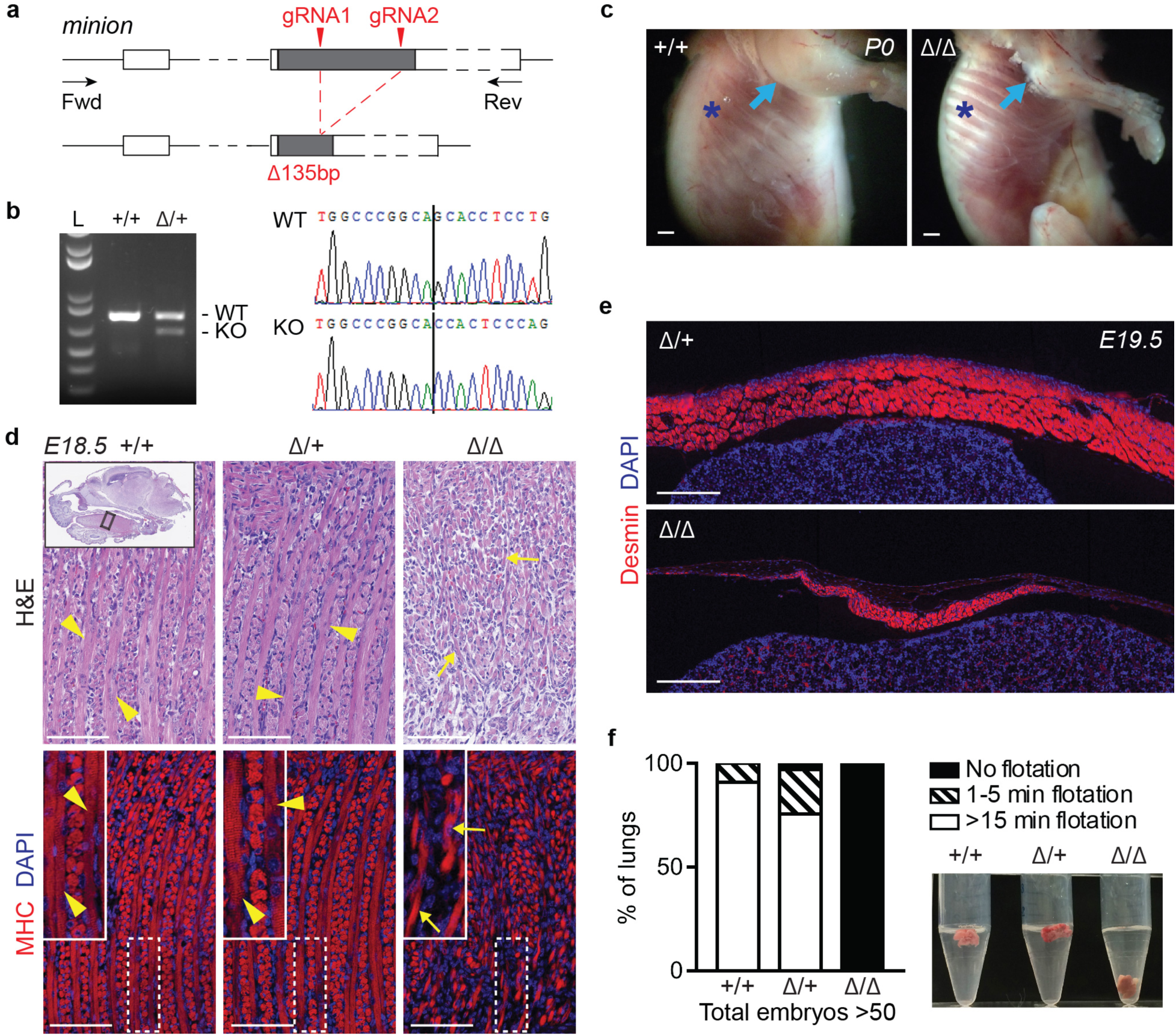
Minion is required for skeletal muscle development. **(a)** Strategy for CRISPR/Cas9 mutagenesis of the *gm7325/minion* locus using a dual sgRNA approach. Gray box, *minion* ORF; white box, non-coding exons; sgRNA, single guide RNA; Fwd and Rev, forward and reverse genotyping primers. **(b)** Left: Representative genotyping PCR of *minion* wild type (+/+) and heterozygous (Δ/+) mice carrying the 135-bp deletion depicted in (a). n=20 (more than 300 adult mice). Right: representative sequence traces. Black line indicates 5’ boundary of the deletion. WT: wild type allele; KO: knockout allele (135-bp deletion). **(c)** Photographs of skinned *minion*^+^*^/^*^+^ and *minion*^Δ/Δ^ P0 mice. Cyan arrows and Blue asterisks indicate forelimb and intercostal musculature, respectively. n=3. **(d)** Histological and immunofluorescence (IF) analyses of embryonic tongue skeletal muscle. Yellow arrowheads and yellow arrows indicate respectively multinuclear myofibers and unfused differentiating elongating myoblasts. Top row: hematoxylin and eosin (H&E) staining of sagittal tongue sections. Inset demonstrates the originating region and orientation of the provided tongue sections. Bottom row: IF staining for the muscle marker MHC (red), with DNA counterstain DAPI (4’,6-diamidino-2-phenylindole; blue). Insets demonstrate magnification of the boxed areas. n=3. **(e)** IF of sagittal sections of diaphragm muscle stained for the muscle marker Desmin (red) and DAPI (blue). n=2 (4 different sections each). **(f)** Quantification (left) and representative image (right) of lung flotation assay using E18.5 mouse embryos (56 total) following 1hr exposure to room air. Scale bars: 1 mm (c), 100 μm (d) and 200 μm (e).

Late stage *minion*^Δ/Δ^ embryos were clearly distinguishable by their decreased size and weight, reduced limb diameter, spinal curvature, and atony, as well as by the dorsal and nuchal subcutaneous edema seen at E16 and earlier stages (Extended Data Fig. 6). *minion*^Δ/Δ^ E17.5 embryos and P0 neonatal pups demonstrated diminutive forelimb and intercostal musculature (Fig. 2c, Extended Data Fig. 6b) and decreased total size of muscle groups, despite no obvious impairment at early embryonic stages (Extended Data Fig. 7). Clear abnormality in skeletal muscle formation was seen at E18.5-19.5, as judged by both histology and immunofluorescence staining for the muscle cell markers MHC and Desmin. Whereas control tongue skeletal muscle contained abundant elongated polynucleated (≥3 nuclei) myotubes, *minion*^Δ/Δ^ muscle demonstrated marked reduction in fused fibers, with accumulation of both short nascent fibers as well as unfused mononucleated cells (Fig. 2d, Extended Data Fig. 8). Similar defects were present in *minion*^Δ/Δ^ forelimb, diaphragm, intercostal, facial, and jaw musculature (Fig.2e, Extended Data Figs. 7d and 9). The observed defects suggested that the perinatal lethality of *minion*^Δ/Δ^ animals could reflect disruption of respiratory function, a possibility we further tested by assessment of lung inflation. Late stage fetuses were delivered by cesarean section and monitored for 1 hour during exposure to room air. Of note, all E18.5 *minion*^Δ/Δ^ embryos died soon after delivery. After 1 hour, lungs were dissected and subjected to flotation testing. In keeping with the dramatic decrease in diaphragm and intercostal muscle formation, lungs from *minion*^Δ/Δ^ but not control animals failed to float (Fig. 2f), demonstrating absence of lung inflation.

The marked reduction of polynucleated myofibers in *minion*^Δ/Δ^ muscle suggested that minion might specifically function in the process of myoblast fusion. Indeed, induction of differentiation in *minion*^Δ/Δ^ primary embryonic myoblasts resulted in near complete failure to form polynucleated myotubes (Fig. 3a, b, Extended Data Fig. 10). Importantly, markers of myogenic commitment and terminal differentiation were induced normally in *minion*^Δ^*^/^*^Δ^ myoblasts both *in vivo* and *in vitro* (Fig. 2d, Extended Data Figs. 8-10), suggesting that the muscle formation defect did not result from a block to progenitor differentiation *per se*. This was further confirmed using loss-of-function in immortalized mouse myoblasts via stable lentiviral transduction with shRNAs targeting the *minion* coding sequence and 3’ untranslated region (UTR) (Extended Data Fig. 11a). Near complete suppression of minion expression was achieved using individual shRNAs (Extended Data Fig. 11b), and a combination of the two most active shRNAs resulted in undetectable protein levels in differentiating cells (*minion*^KD^; Fig. 3c). Analysis of Myogenin, MyoD, Desmin, and MHC expression confirmed both the absence of any molecular differentiation defect in *minion*^KD^ cells (Fig. 3c, e, Extended Data Fig. 11c), as well as the presence of a severe block to myoblast fusion (Fig. 3d, f, g). Interestingly, differentiating *minion*-deficient myoblasts elongated and aligned normally despite failing to fuse (Fig. 3d), suggesting that myoblast apposition was not impaired. Similar results were obtained using lentiviral shRNA transduction of primary, non-immortalized adult mouse myoblasts (Extended Data Fig. 12).

**Fig. 3.**
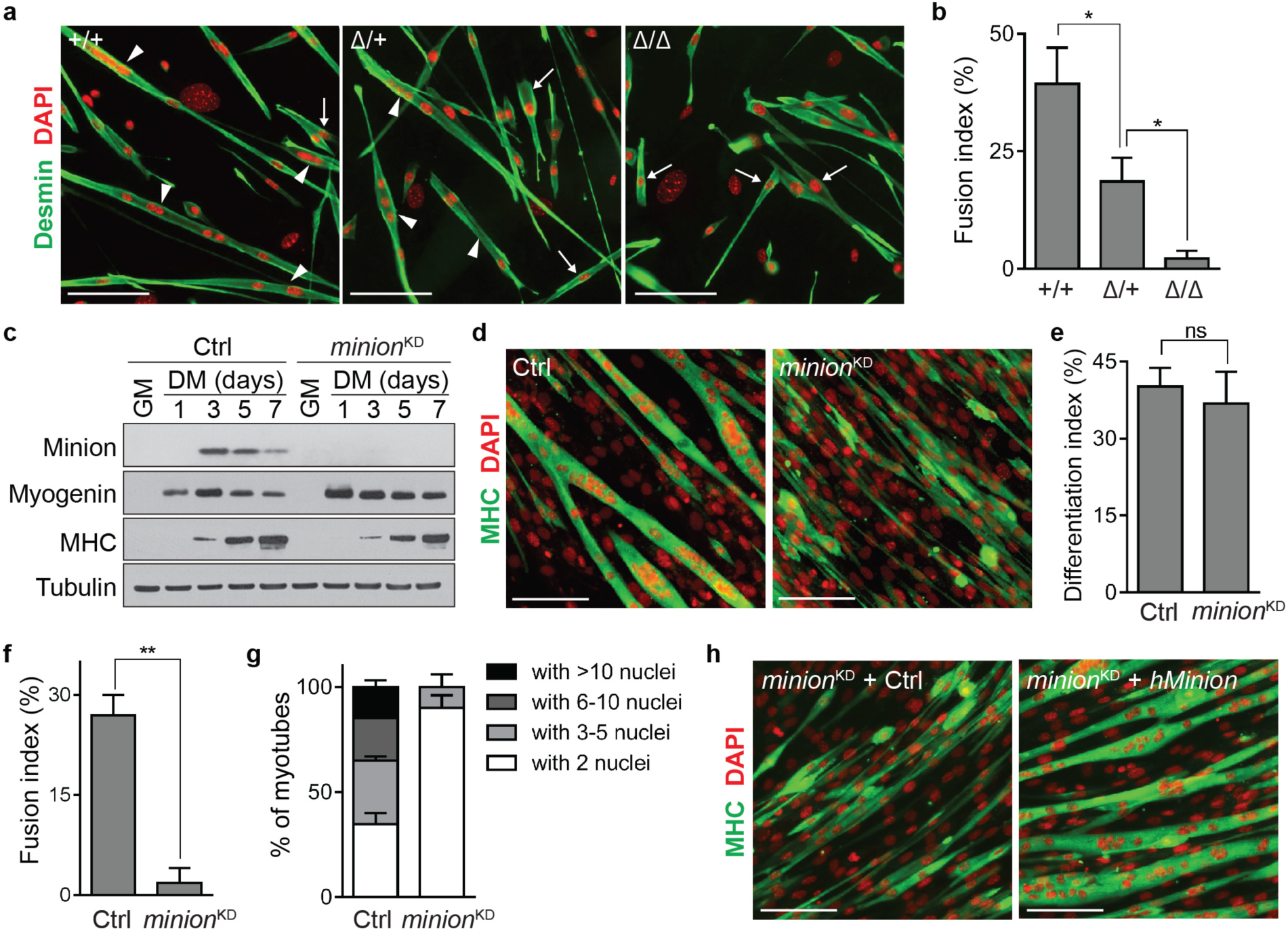
minion is specifically required for fusion of skeletal muscle progenitors. **(a)** Immunofluorescence of primary embryonic myoblasts isolated from E18.5 *minion*^+^*^/^*^+^, *Minion* ^Δ^*^/^*^+^ and *minion* ^Δ/Δ^ embryos, following 3 days in DM. Desmin (green) and DAPI (red). White arrowheads: myotubes; white arrows: elongating myoblasts. n=3 (5 technical replicates each). **(b)** Fusion index of myoblasts in (a), calculated as % nuclei in Desmin^+^ myotubes (≥ 3 nuclei) of total nuclei in Desmin^+^ cells. Asterisk: *P* <0.05. n=4 (two 0.7mm×7mm fields each). **(c)** Western blots of C2C12 myoblasts cultured in GM or DM. Cells were lentivirally infected with either control *luciferase* targeting shRNA (Ctrl) or serially with two shRNA targeting the *minion* 3’UTR (*minion*^KD^) and cultured in GM or DM for the indicated number of days. n=3. **(d)** IF of Ctrl and minion^KD^ myofibers following 5 days in DM. MHC (green) and DAPI (red). n=5 (8 technical replicates each). **(e)** Differentiation index for (d), calculated as % nuclei in MHC^+^ cells of total nuclei. NS: not significant. **(f)** Fusion index for (d), calculated as % nuclei in MHC^+^ myotubes (≥ 3 nuclei) of total nuclei. Double asterisks: *P* <0.001. **(g)** Quantification of myotubes by nuclei number for (d). e-g, n=4 (two 0.7mm×0.7mm fields each). **(h)** IF of minion^KD^ cells expressing either control protein (NanoLuc) or human Minion ortholog, after 5 days in DM. MHC (green) and DAPI (red). n=3 (6 technical replicates each). Scale bars: 100 μm.

As the shRNAs used to target the *minion* transcript in *minion*^KD^ cells recognize the 3’ UTR, we tested the ability of various ORF cDNA clones to complement the *minion*^KD^ cell fusion defect (Extended Data Fig. 13a). Both full-length untagged and C-terminally tagged mouse minion robustly rescued myoblast fusion (Extended Data Fig. 13b), demonstrating that the fusion defect observed in *minion*^KD^ cells was not the result of off-target effects. The putative human Minion ortholog, previously annotated as a long noncoding RNA (GRCh37 genome assembly), was then tested in a similar complementation assay, demonstrating that both untagged and C-terminally epitope-tagged human *Minion* ORFs strongly reconstituted cell fusion in *minion*^KD^ cells (Fig. 3h, Extended Data Fig. 13b). To definitively establish that these ORFs function via protein coding, single nucleotide insertions or deletions were introduced into the untagged mouse and human *Minion* cDNAs, respectively. These frameshift point mutants failed to complement the fusion defect (Extended Data Fig. 14), confirming that these transcripts function not as non-coding RNAs but by encoding functional microproteins. Reconstitution of *minion*^KD^ cells with cDNA mimicking the 135bp-deletion allele found in the *minion*^Δ/Δ^ mice likewise failed to rescue cell fusion (Extended Data Fig. 15), confirming that this represents a true loss-of-function allele. Taken together, these data demonstrate that *minion* encodes a microprotein essential for skeletal muscle formation via a specific function in myoblast fusion.

The requirement for minion in cell fusion appears muscle specific, as minion expression was not seen in other settings of physiologic cell fusion, such as in the placenta or in fusing macrophage lineage cells (Extended Data Fig. 16). This restricted expression pattern mirrors that of myomaker, a recently described transmembrane regulator of myoblast fusion, and we therefore investigated the functional relationship between these proteins in order to better understand the mechanism of minion-associated cell fusion. Myomaker was readily detectable in regenerating muscle and differentiating mouse myoblasts (Extended Data Fig. 17), when minion levels are likewise high (Fig. 1b, d). Loss of minion expression did not impair myomaker expression in differentiating myoblasts (Fig. 4a), and myomaker overexpression was incapable of rescuing the fusion defect in *minion*^KD^ cells (Fig. 4b, c). We therefore conclude that the cell fusion defect seen in the absence of minion function does not reflect deficiency in myomaker expression. Co-immunoprecipitation revealed no detectable physical interaction between the two proteins in differentiating muscle cells (data not shown).

**Fig. 4.**
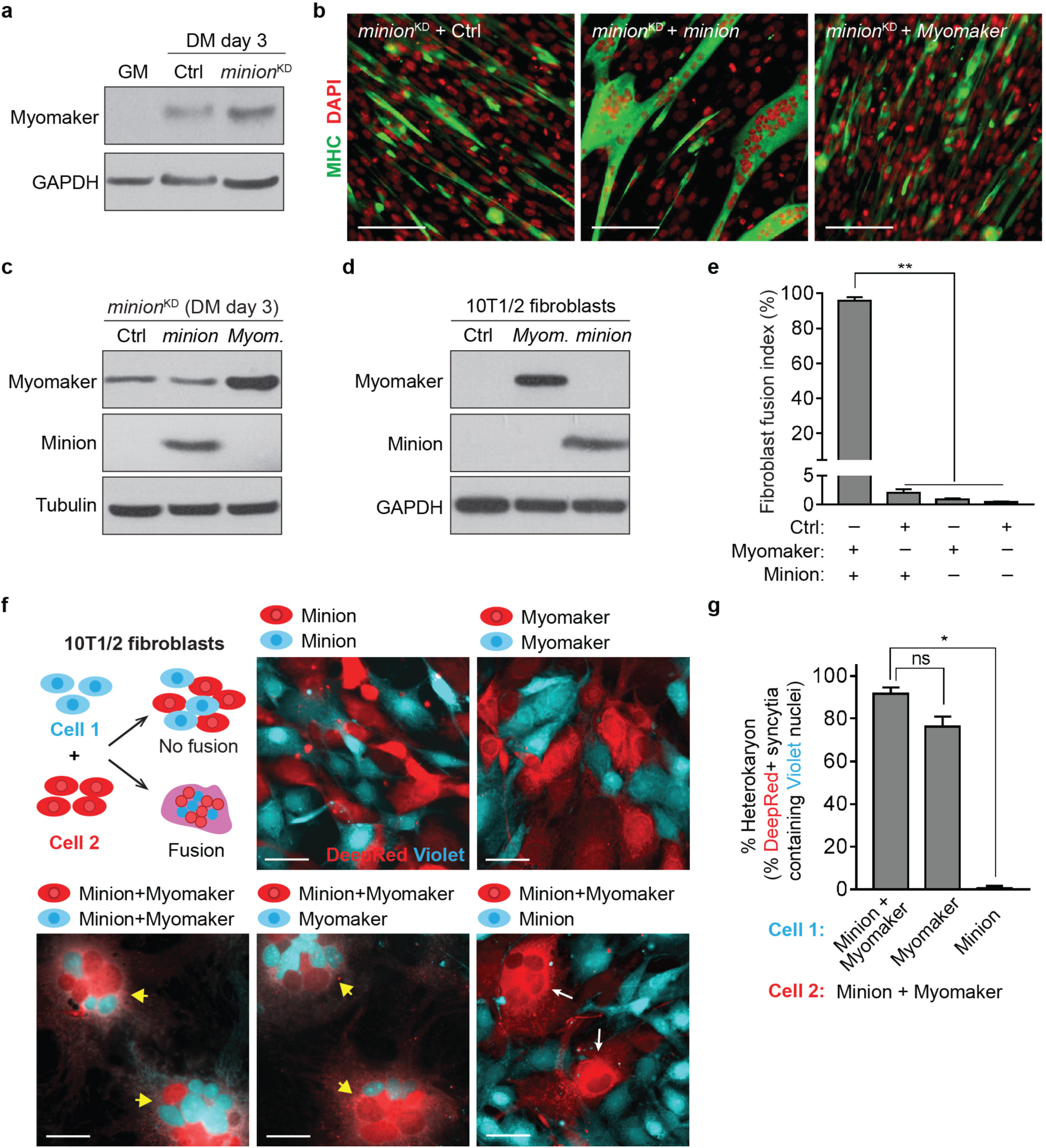
Coexpression of minion and myomaker is sufficient to induce cell fusion in a heterologous system. **(a)** Western blot of wild-type C2C12 myoblasts in GM, as well as Ctrl and *minion*^KD^ myoblasts in DM for 3 days. n=2. **(b)** IF of Ctrl and *minion*^KD^ myoblasts expressing Luciferase (Ctrl), minion, or myomaker, after 5 days in DM. MHC (green) and DAPI (red). n=2. **(c)** Western blot of cell lines shown in (b). n=2. **(d)** Western blot of 10T1/2 fibroblasts expressing Luciferase (Ctrl), myomaker, or minion. n=2. **(e)** Quantification of GFP^+^ syncytia (≥3 nuclei) in fibroblasts expressing combinations of proteins as indicated. Syncytia were scored 24hr after seeding. n=3 (two 1.4mm×1.4mm fields each). **(f)** Fluorescence images from cell mixing experiments using differentially labeled 10T1/2 fibroblasts. Cells were serially infected with retroviruses encoding the indicated combinations of minion, myomaker, or controls (label omitted for simplicity). CellTrace Violet (blue) and CellTracker DeepRed (red) dyes were used for labeling. Yellow arrowheads indicate syncytia containing both DeepRed^+^ cells and Violet^+^ nuclei. White arrows indicate syncytia derived from DeepRed^+^ cells only. n=5 (6 technical replicates each). **(g)** Quantification of fusion in (f) (bottom panels), measured as percentage of DeepRed^+^ syncytia (≥ 3 nuclei) containing ≥ 1 Violet^+^ nucleus. n=4 (six 1.4mm×1.4mm fields each). Scale bars: 100 μm **(c)**, and 50 μm **(f).**

Previous studies have demonstrated that expression of myomaker alone fails to induce fusion between non-myogenic cells, and that at least one additional, as yet unidentified factor is required^4^. We likewise observed that heterologous expression of neither myomaker nor minion alone in 10T1/2 fibroblasts was sufficient to drive fusion of these cells with one another (Fig. 4d, f, Extended Data Fig. 18, and data not shown). Remarkably however, we observed that simultaneous expression of minion and myomaker together drove rapid and uniform fusion of transduced fibroblasts with one another, leading to the formation of large multinuclear syncytia (Fig. 4e and Extended Data Fig. 18). Similar results were observed in undifferentiated myoblasts cultured under growth conditions (data not shown).

Mechanistically, the small size and lack of functional domains within microproteins has led to the suggestion that they function primarily via protein-protein interactions^1^. We therefore performed affinity purification followed by mass spectrometry analysis using FLAG-tagged minion expressed in differentiating myoblasts. This identified several classes of highly enriched interacting proteins (Extended Data Fig. 19a, b; Extended Data Table 1), including cytoskeletal proteins. Indeed, we observed that co-expression of minion and myomaker in fibroblasts induced dramatic cytoskeletal rearrangement, with formation of an actin wall at the cell periphery^16^ (Extended Data Fig. 19c). Treatment with two different actin-polymerization inhibitors, which disrupt cytoskeleton remodeling, blocked both actin reorganization and cell fusion in this minimal two-factor system (Extended Data Fig. 19d). To further demonstrate that syncytium formation induced by minion and myomaker represents true cell fusion rather than incomplete cytokinesis, cell mixing experiments were performed using fluorescently labeled populations expressing either minion, myomaker, or both proteins (Fig. 4f, g, Extended Data Fig. 20). In addition to confirming rapid cell fusion, an unexpected but clear polarity was observable within the fusion pair, with minion expression required in only one cell, whereas myomaker expression was required within both fusing cells (Fig. 4f, g, Extended Data Fig. 20). We conclude that minion is the previously unknown factor required for myomaker to mediate fusion of cells into differentiating skeletal muscle, and that minion and myomaker can together function as a minimal program for the induction of cytoskeletal rearrangements leading to fusion in otherwise non-fusogenic cells.

The data presented here uncover an evolutionarily conserved pathway for cell fusion mediated by the microprotein minion and the transmembrane protein myomaker. Our studies reveal an unanticipated polarity within the fusion pair, in which both cells must express myomaker but only one cell need express minion in order to drive cell fusion. This suggests that vertebrate muscle formation has previously unrecognized similarities with invertebrate muscle development, in which fusion requires active cytoskeletal remodeling and occurs between distinct populations of founder cells and fusion competent myoblasts^16-22^. We suggest a model in which the transmembrane protein myomaker induces apposition or adhesion of cell membranes^23^, whereas the microprotein minion drives the cytoskeletal reorganization needed for fusion. Moreover the two-factor fusion system described here opens the door to programmable and potentially targeted cell fusion, which may find therapeutic and research application in oncolytic fusion of cancer cells^24^, fusion of dendritic cells to cancer cells in immunotherapy^25^, therapeutic fusion in regenerative medicine^26,27^, and heterokaryon-based studies of nuclear reprogramming^28^. Finally and importantly, our studies constitute the first report, to our knowledge, of an essential mammalian microprotein. This represents the strongest evidence to date that this class of diminutive proteins in fact constitutes a ‘microproteome’ with critical and largely unexplored functions.

## Acknowledgements

The authors gratefully acknowledge Qiang Zhou, Cynthia Cienfuegos, and Jacqueline Avis for reverse genetics support; Brian Schwartz and Robbin Newlin for histology support; Fabio Luna, Whitney Barnes, and John Walker for RNA sequencing; Doug Quackenbush for assistance with microscopy; Minhua Qiu for image analysis; Albert Parker for advice regarding molecular biology; Daniel Mason for assistance with mass spectrometry; Seung-Hyun Woo for advice regarding lung flotation assay; Jason Matzen, Erik Spedale, and Paul Calvin for assistance with compounds; Bin Fang for bioinformatic assistance; and Melanie Tucker, Mary Frazer, Diego Guzman, and Richard Eddins for animal husbandry support. We thank David Glass, Richard Glynne, and Peter McNamara for leadership support, and David Glass, Estelle Trifilieff, Richard Glynne, Peter McNamara, John Walker, and Rajesh Nair for critical reading of the manuscript.

---

**Extended Data Fig. 1.**
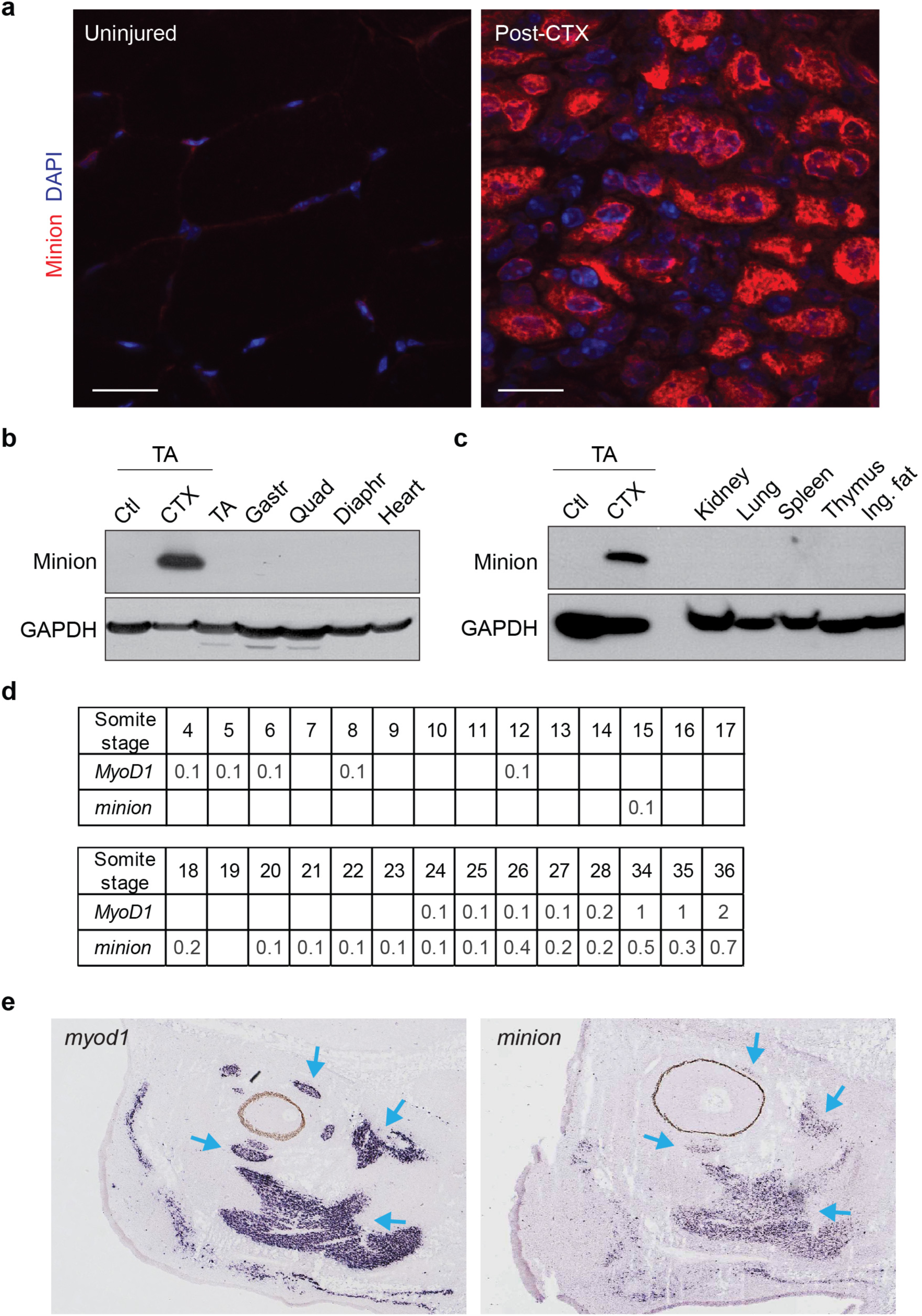
Minion is highly and specifically expressed in regenerating and developing skeletal muscle. **(a)** Immunofluorescence staining of transverse sections prepared from fresh frozen uninjured (left) or regenerating (right, 3 to 4 days post CTX injection) TA muscle. minion (red) and DAPI (blue) are shown. Scale bar: 20 μm. n=2 (5 different sections each). **(b)** Western blot analysis of various adult skeletal muscle groups and of cardiac muscle using the indicated antibodies. 6-8 weeks old C57BL/6J mice were used. Ctl: uninjured TA muscle; CTX: regenerating TA muscle at day 4 post CTX injection; TA: *tibialis anterior;* Gastr: *gastrocnemius;* Quad: *quadriceps femoris;* Diaphr: diaphragm. n=3. **(c)** Western blots analysis using the indicated antibodies of various adult tissues from 6-8 week C57BL/6J mice. n=3. **(d)** Expression of mouse *minion* during embryonic development as detected by RNA-seq (EMBL-EBI Expression Atlas; http://www.ebi.ac.uk/gxa)^29,30^. Numbers indicate RNA expression level (FPKM). **(e)** Expression of mouse *myod1* (left) and *minion* (right) in non-somitic muscle as detected by *in situ* hybridization at E14.5 (Eurexpress; http://www.eurexpress.org/ee/intro.html)^31^. Blue arrows indicate overlapping expression in extraocular and facial muscle.

**Extended Data Fig. 2.**
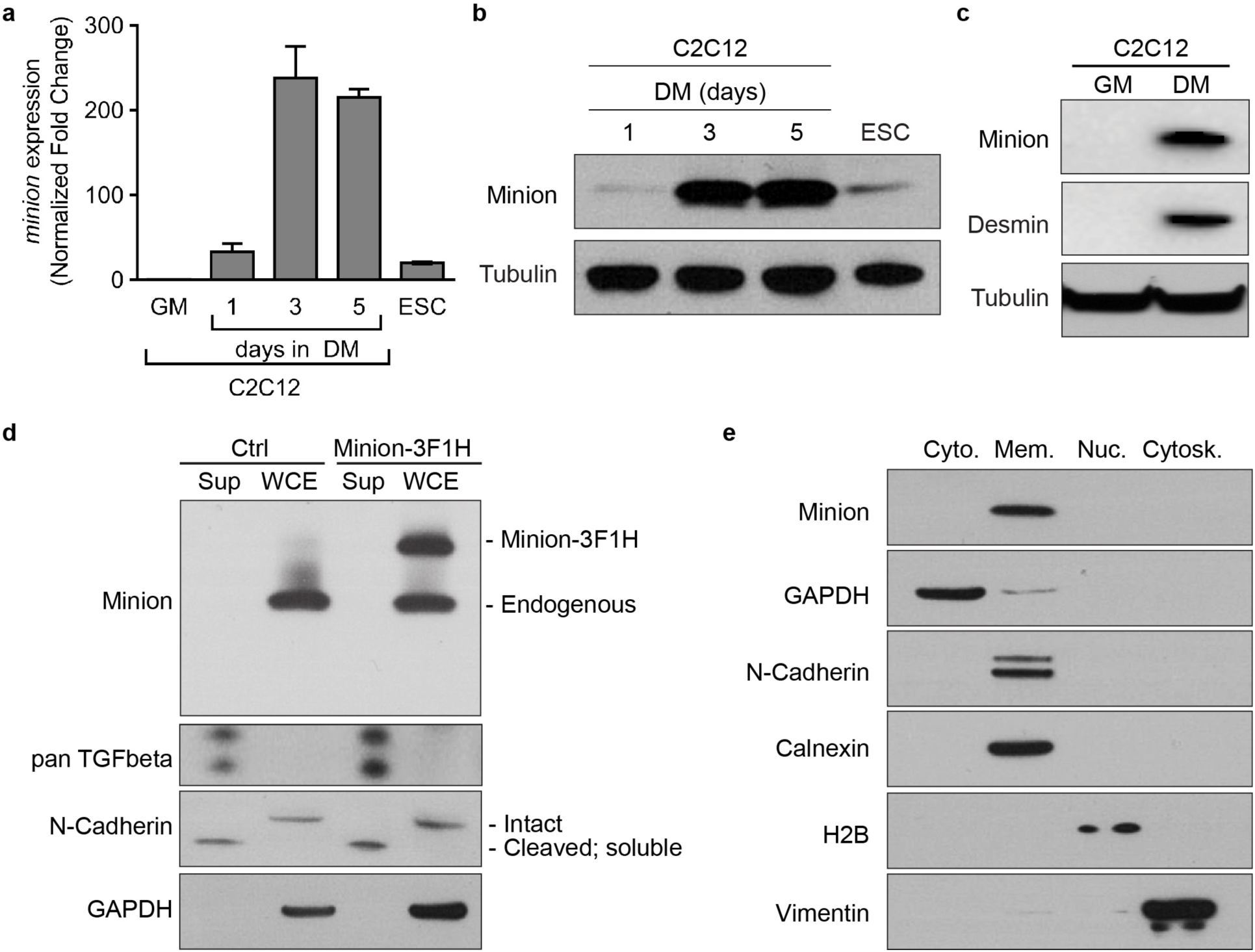
*minion* is highly expressed in differentiating muscle cells and is membrane associated. **(a)** RT-qPCR quantification of *minion* mRNA levels in C2C12 myoblasts under growth (GM) or differentiation (DM) conditions. Mouse embryonic stem cells (ESC) served as a positive control. n=3. **(b)** Western blot analysis of differentiating C2C12 myoblasts (at day1, 3, 5 in DM), and of embryonic stem cell (ESC) using the indicated antibodies. n=2. **(c)** Western blots analysis of C2C12 myoblasts in GM or at day3 in DM using the indicated antibodies. Desmin is an intermediate filament protein high in differentiated muscle cells. n=2. **(d)** Western blot analysis of concentrated cell culture supernatant (Sup) from day 3 differentiating C2C12 myoblasts expressing control vector or C-terminally 3×FLAG-1×HA-tagged Minion. Both tagged and endogenous Minion are detected in whole cell extract (WCE). TGFβ and cleaved N-Cadherin are positive controls, whereas GAPDH and intact N-Cadherin are non-secreted negative controls. n=3. **(e)** Subcellular fractionation of C2C12 myoblasts at day 4 in DM. Western blot analysis was performed using the indicated antibodies. The four fractions examined were cytosolic (Cyto), membrane (Mem, nuclear (Nuc), and cytoskeletal (Cytosk). The membrane fraction contains both plasma membrane as well as intracellular membranes (ER, Golgi, mitochondria, endosome, lysosome, etc.). GAPDH: cytosolic fraction marker; N-Cadherin: plasma membrane marker; Calnexin: ER membrane marker; Histone H2B: nuclear fraction marker; Vimentin: cytoskeletal fraction marker. n=3.

**Extended Data Fig. 3.**
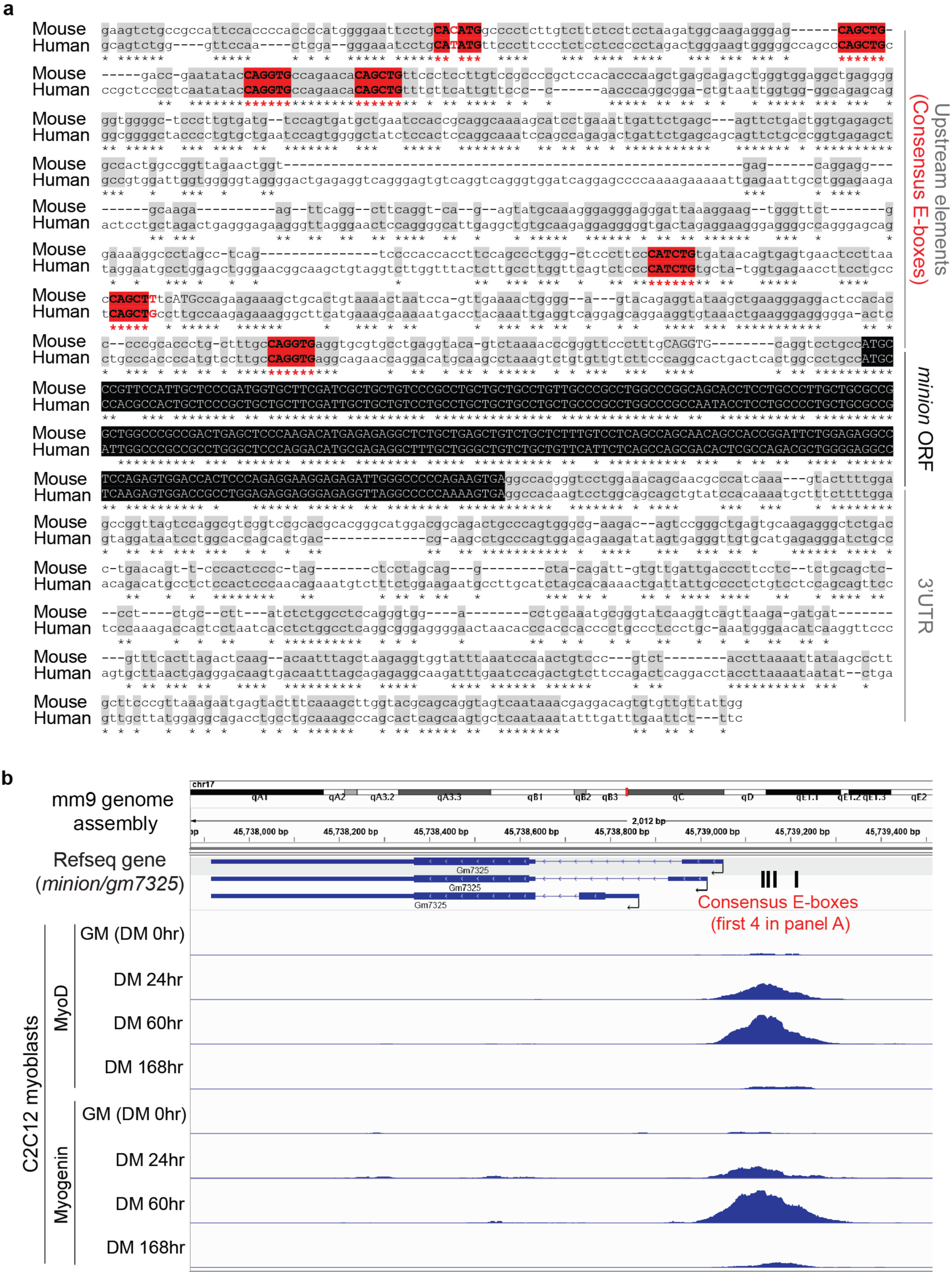
*minion* promoter structure is consistent with regulation by the muscle regulatory factors MyoD and Myogenin. **(a)** Sequence alignment of the genomic regions surrounding mouse *gm7325/minion* gene and its human ortholog *RP1-302G2.5* (LOC101929726). The *minion* ORF is highlighted in black. Seven conserved E-box motifs (CANNTG and CANNTT) within the promoter and 5’UTR region are highlighted in red. **(b)** ENCODE MyoD and Myogenin ChIP-seq data from C2C12 myoblasts under growth conditions (GM) and at three time points under differentiation conditions (DM) were examined surrounding the *minion* genomic locus. Black bars indicate the first four conserved E-box motifs within the promoter region in (a).

**Extended Data Fig. 4.**
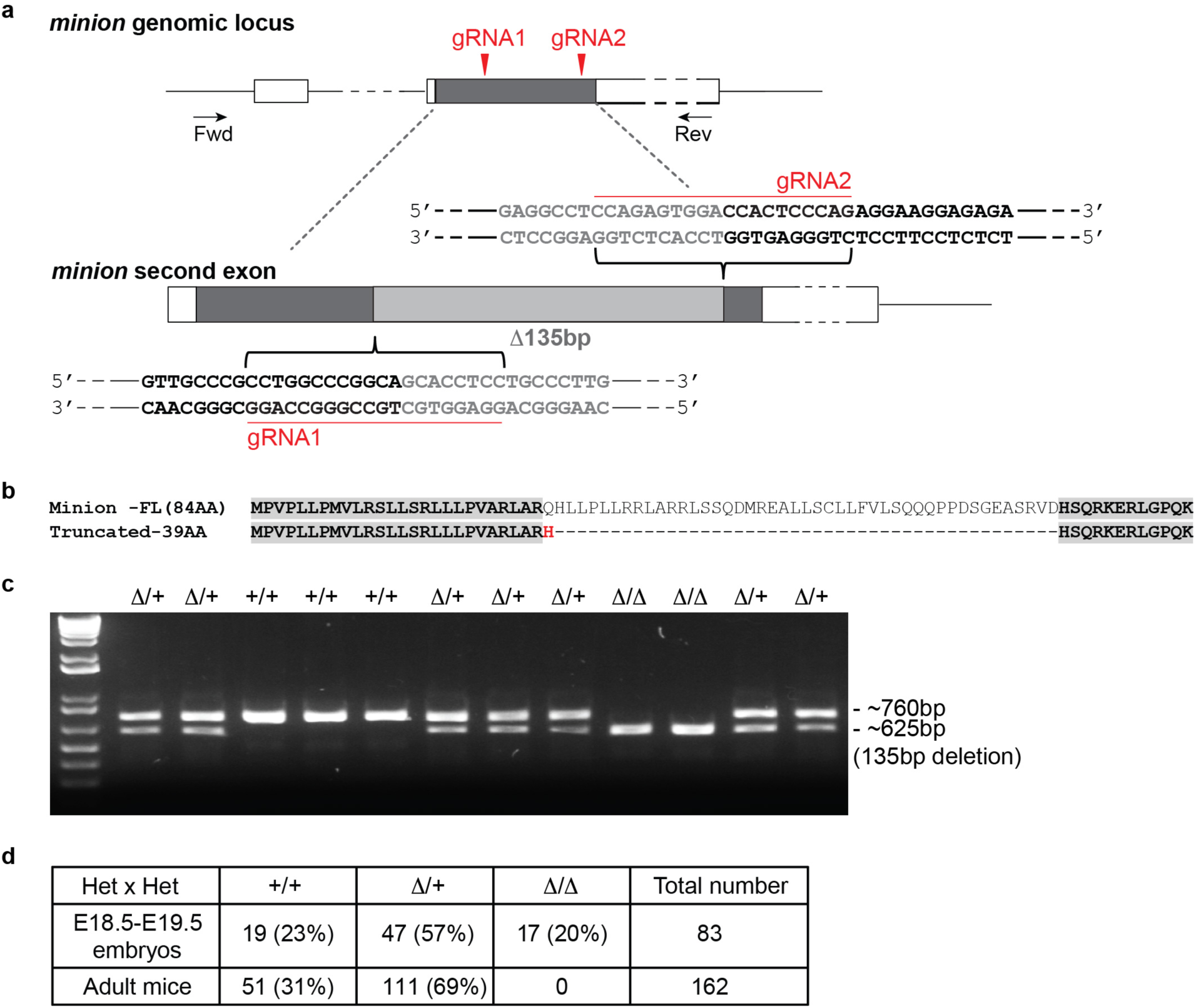
Generation of *minion*-deficient knockout mice by CRISPR/Cas9-mediated genome editing. **(a)** Detailed schematic of Fig.2A for generation of *minion*-deficient animals by CRISPR/Cas9 mediated editing. A 135bp in-frame deletion (marked in light gray) within the *gm7325/minion* ORF (gray) was induced using a double sgRNA approach. The genomic target sequences of the gRNAs are indicated with a red line. Fwd: forward PCR genotyping primer; Rev: reverse PCR genotyping primer. **(b)** Protein sequence alignment of the 84 aa full-length mouse minion and the predicted 39 aa truncated form (minion ^Δ^). The truncated form is predicted to contain only the N-terminal 26 aa and C-terminal 12 aa of the original microprotein. **(c)** Agarose gel picture of typical genotyping PCR results using E18.5 embryos from *minion*^Δ/+^*× minion* ^Δ/+^ crosses (Het×Het). +/+: wild type (*minion*^+/+^); Δ/+: heterozygote (*minion*^Δ/+^); Δ/Δ: *minion* knockout homozygote (*minion*^Δ/Δ^). n=12 (83 E18.5-E19.5 embryos) **(d)** Summary of genotyping results of both late-stage embryos and adult mice from *minion*^Δ/+^ × *mimon*^Δ/+^ crosses.

**Extended Data Fig. 5.**
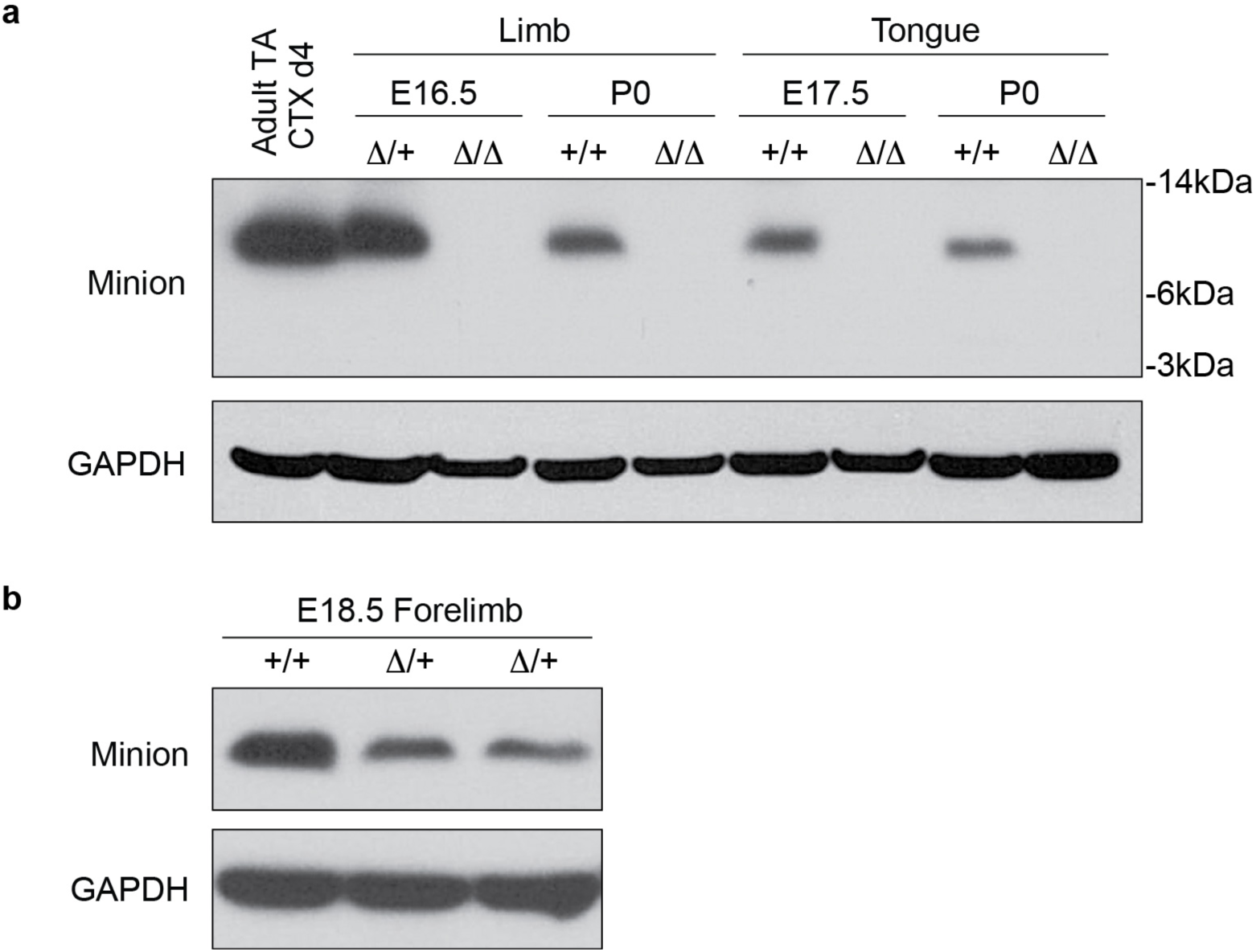
Loss of minion protein expression in *minion*-deficient animals. **(a)** Western blot analysis of limbs and tongues from E16.5/E17.5 embryos and P0 mice using indicated antibodies. Adult TA muscle at 4 days post cardiotoxin injection (CTX 4dpi) was used as a positive control. Embryos from the same litter were used for each comparison. n=3. The full-length minion was not observed in the *minion* ^Δ/Δ^ embryos, and the predicted 39 aa truncated protein was not observed either using the same anti-minion antibody. **(b)** Western blot analysis of forelimbs from *minion*^+/+^ and *minion* ^Δ/+^ E18.5 embryos of the same litter using indicated antibodies.

**Extended Data Fig. 6.**
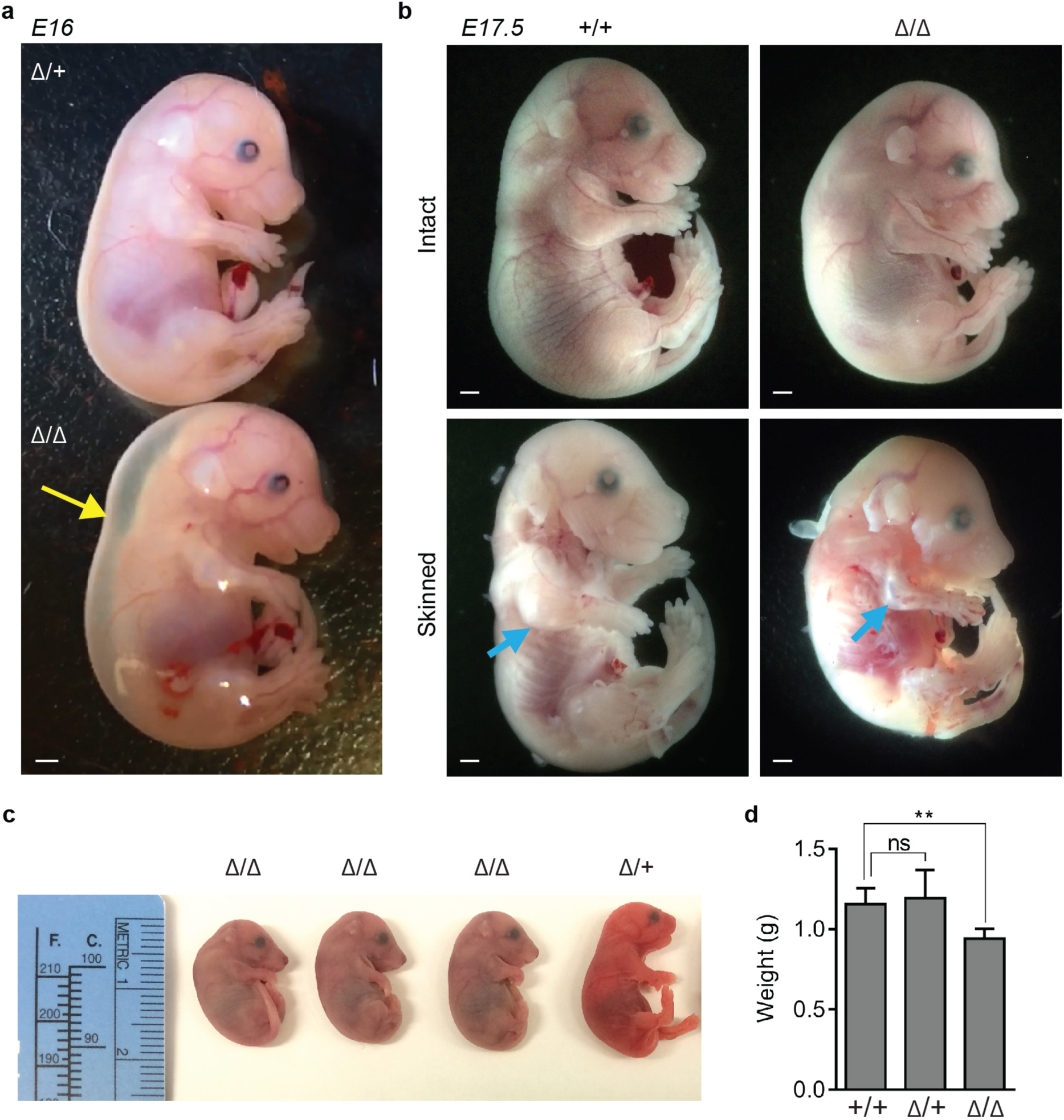
Developmental abnormalities in *minion*-deficient animals. **(a)** Photos of *minion* ^Δ/+^ and *minion*^Δ/Δ^ E16 embryos. Yellow arrow indicates accretion of dorsal and nuchal subcutaneous edema. n=3. Scale bar: 1 mm. **(b)** Photos of *minion*^+^*^/^*^+^ (left) and *minion* ^Δ/Δ^ (right) E17.5, either unskinned (top) or skinned (bottom). Cyan arrow indicates expected location of forelimb musculature. n=3. Scale bar: 1 mm. **(c)** Photos of E18.5 embryos of the indicated genotypes following air breathing after delivery by cesarean section. The *minion* ^Δ/Δ^ embryos were atonic and exhibited an abnormal spinal curvature and became cyanotic and dead soon after delivery. n=5. **(d)** Quantification of E18.5 embryo weight after delivery by cesarean section (33 embryos total). NS: not significant; double asterisks: *P* <*0.001*.

**Extended Data Fig. 7.**
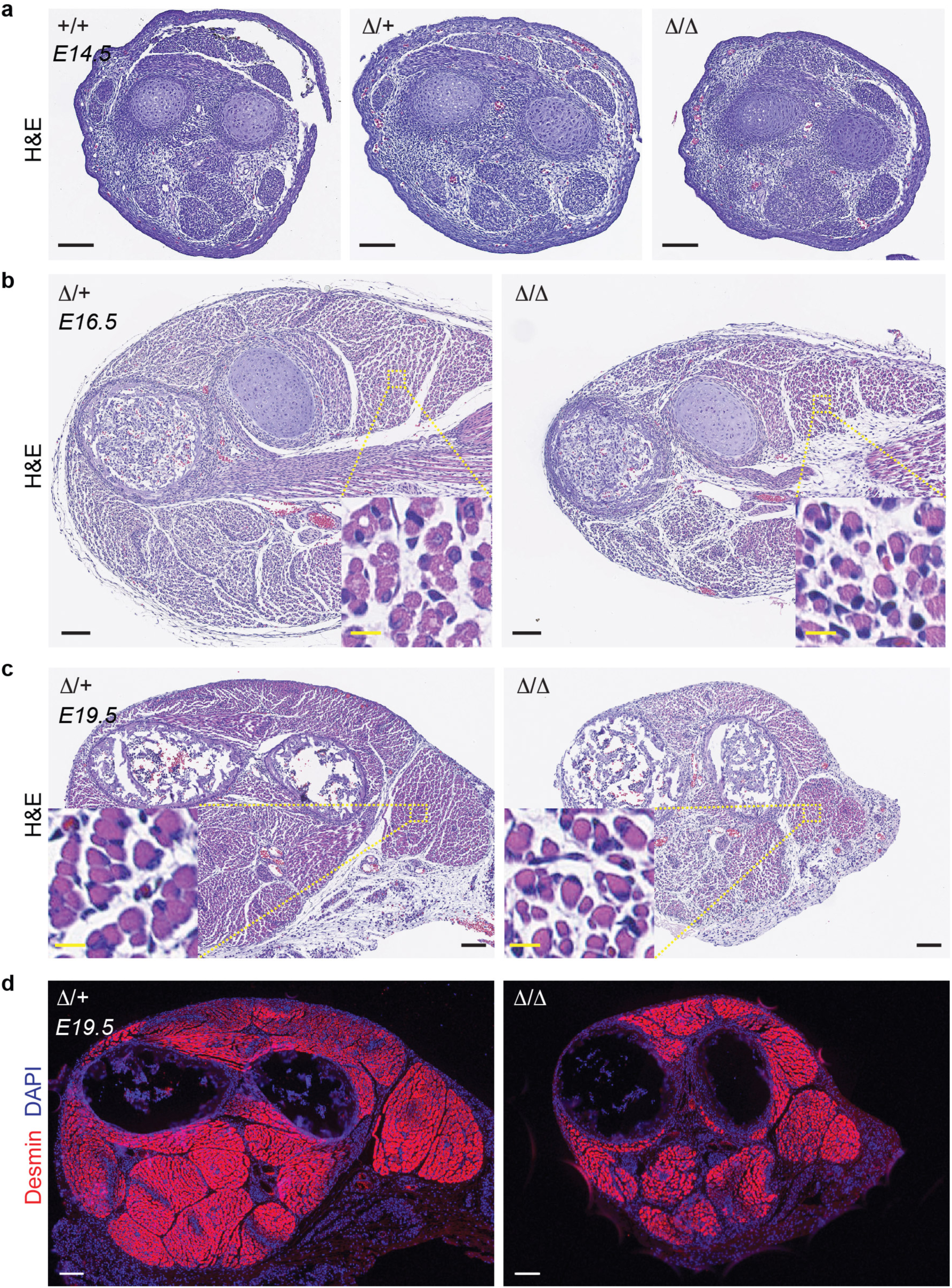
Minion deficiency affects later stages of skeletal muscle formation. Paraffin-embedded embryos of different stages were examined. **(a)** Histological images of hematoxylin and eosin (H&E) stained forelimb transverse sections of E14.5 embryos with indicated genotypes. No significant difference was observed between genotypes with respect to muscle group number, position or size. n=2. **(b)** Histological images of H&E-stained E16.5 forelimb transverse sections of indicated genotypes. Inset demonstrates magnification of the region shown in yellow box. n=2. **(c)** Histological images of H&E-stained forelimb transverse sections for E19.5 embryos with indicated genotypes. Inset demonstrates magnification of the region shown in yellow box. n=3. **(d)** Immunofluorescence images of forelimb transverse sections for E19.5 embryos with indicated genotypes. Desmin (red) marks all differentiating myoblasts, myotubes, and muscle fibers in the embryo. Loss of minion does not obviously block the differentiation of skeletal muscle during development. n=3. Nuclei are labeled by DAPI (blue). Black and white scale bars: 100 μm; yellow scale bars: 10 μm.

**Extended Data Fig. 8.**
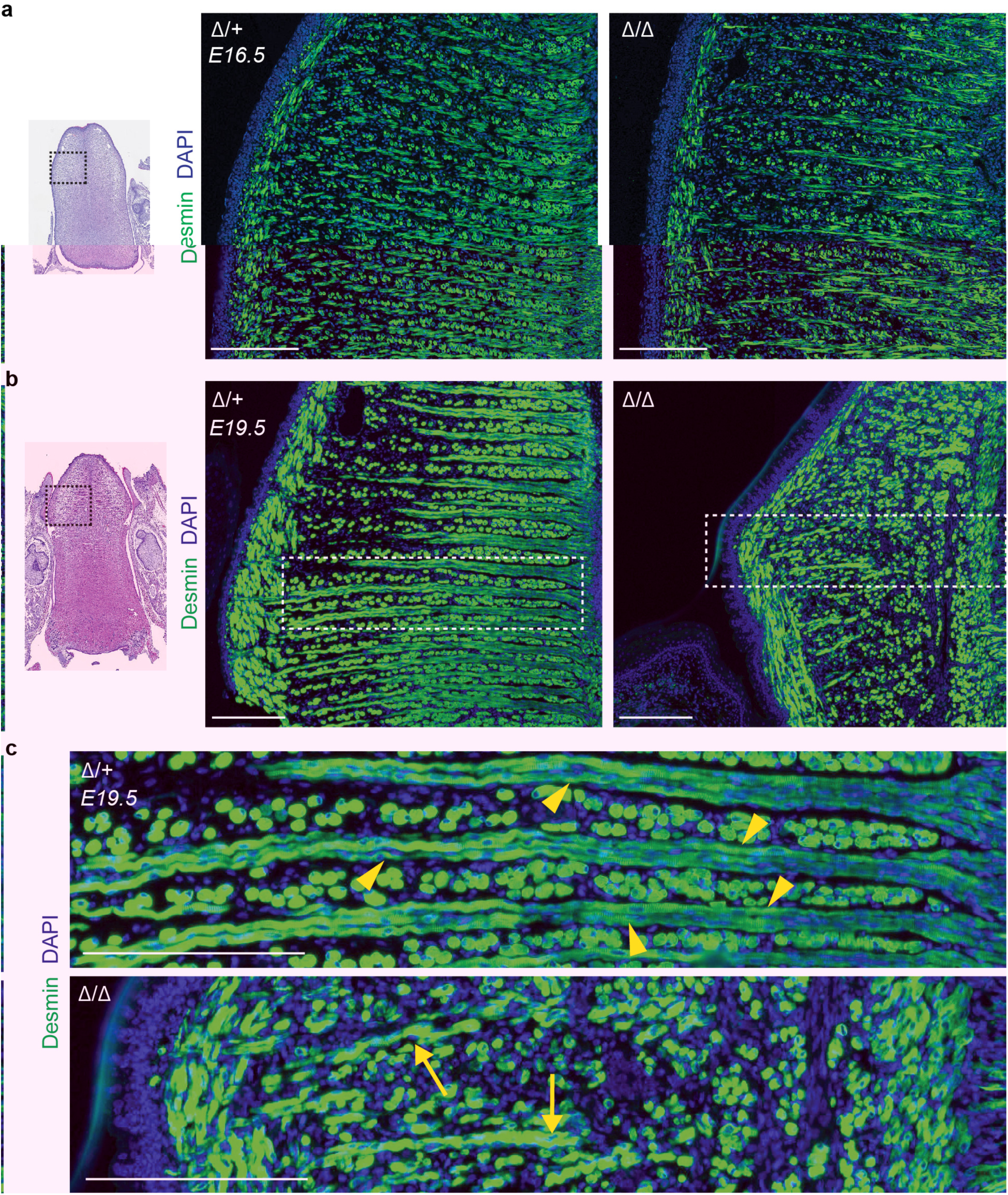
Loss of minion does not affect differentiation of skeletal muscle progenitors *in vivo* but blocks formation of polynucleated fibers. Paraffin-embedded embryonic tongues of different stages were used to examine both longitudinal and transverse myofibers. **(a)** Immunofluorescence images of E16.5 tongue transverse sections. Black box in histological image at left demonstrates the area shown at right in fluorescence images. n=2. Desmin (green) and DAPI (blue) are shown. **(b)** Immunofluorescence images of E19.5 tongue transverse sections. Black box in histological image at left demonstrates the area shown at right in fluorescence images. n=3. White box indicates area magnified in (c). Desmin (green) and DAPI (blue) are shown. **(c)** Magnified view of white boxed area shown in (b), demonstrating significant reduction in polynucleated myofibers in *minion* ^Δ/Δ^ tongue. Yellow arrowheads and yellow arrows indicate fused multinuclear myofibers and unfused differentiating and elongating myoblasts respectively. Scale bars: 200 μm.

**Extended Data Fig. 9.**
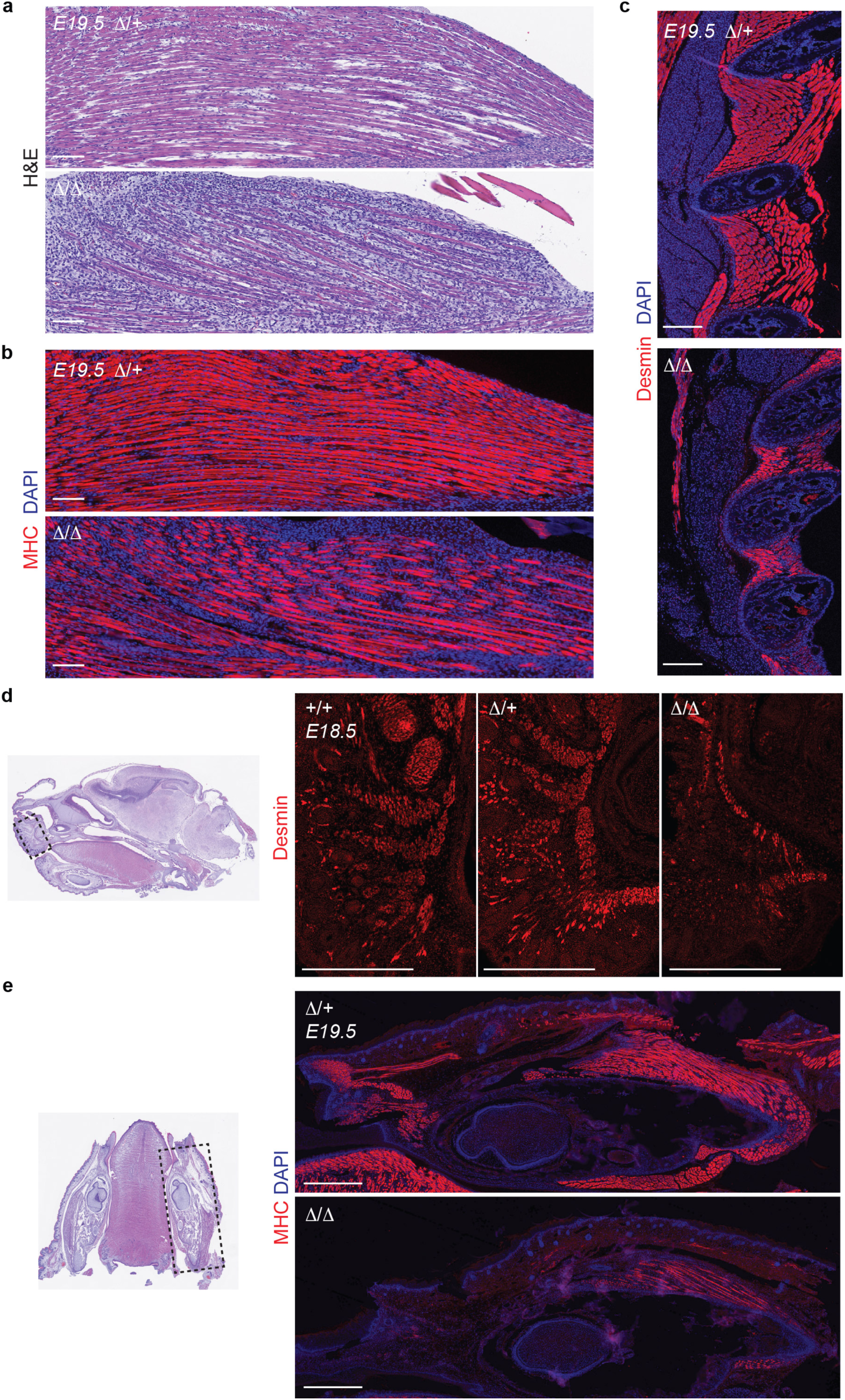
Minion deficiency leads to defects in skeletal muscle formation. Paraffin-embedded embryos of different stages were examined. **(a)** Histological images of H&E-stained E19.5 forelimb longitudinal sections of indicated genotypes. n=3. **(b)** Immunofluorescence images of forelimb longitudinal sections for E19.5 embryos with indicated genotypes. MHC (red) and DAPI (blue) staining are shown. n=3. **(c)** Immunofluorescence images of intercostal muscle sagittal sections for E19.5 embryos with indicated genotypes. Desmin (red) and DAPI (blue) staining are shown. n=2. **(d)** Immunofluorescence images of non-somitic facial musculature sagittal sections from E18.5 embryos with the indicated genotypes. Desmin (red) staining is shown. Black box in histological image at left demonstrates the area shown at right in fluorescence images. n=2. **(e)** Immunofluorescence images of non-somitic branchial (jaw) and facial musculature on transverse sections from E19.5 embryos with the indicated genotypes. n=2. MHC (red) and DAPI (blue) staining are shown. Scale bars: (a-c) 100μm and (d-e) 500μm.

**Extended Data Fig. 10.**
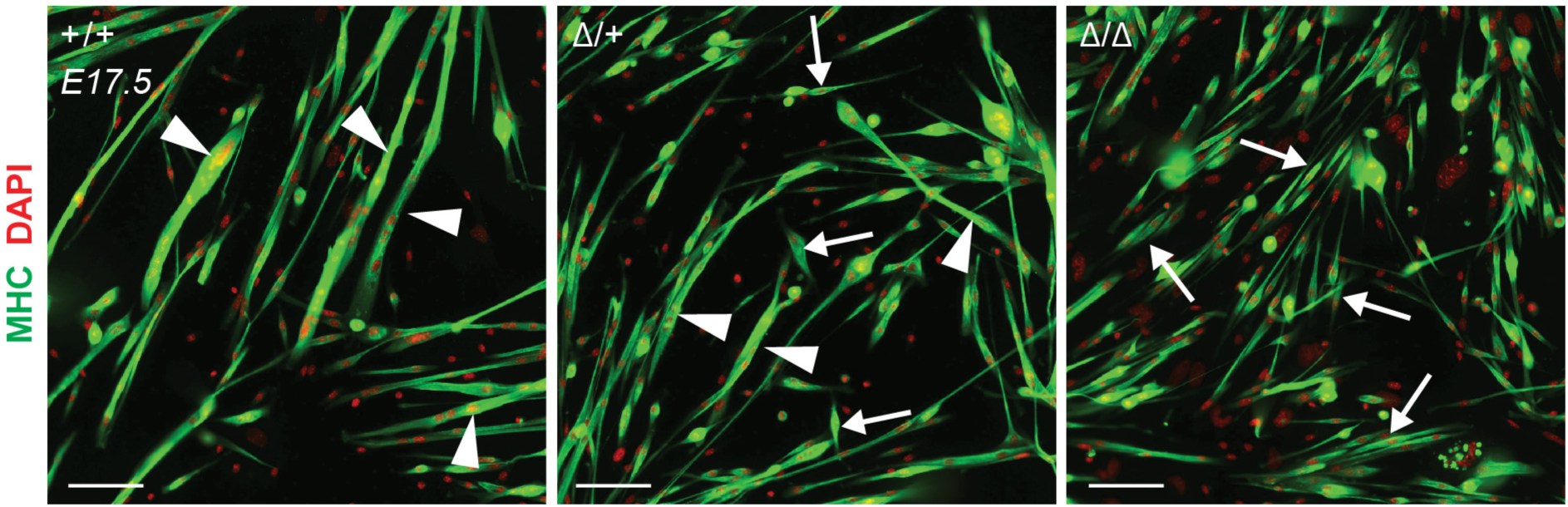
Genetic loss of minion blocks embryonic myoblast fusion *in vitro*. Immunofluorescence images of E17.5 primary embryonic myoblasts isolated from limbs of indicated genotypes following 3 days in differentiation medium. White arrowheads and white arrows point out the fused multinuclear myotubes and unfused differentiating elongating myoblasts. n=2 (5 technical replicates each). MHC (green) and DAPI (red) are shown. Scale bar: 100 μm.

**Extended Data Fig. 11.**
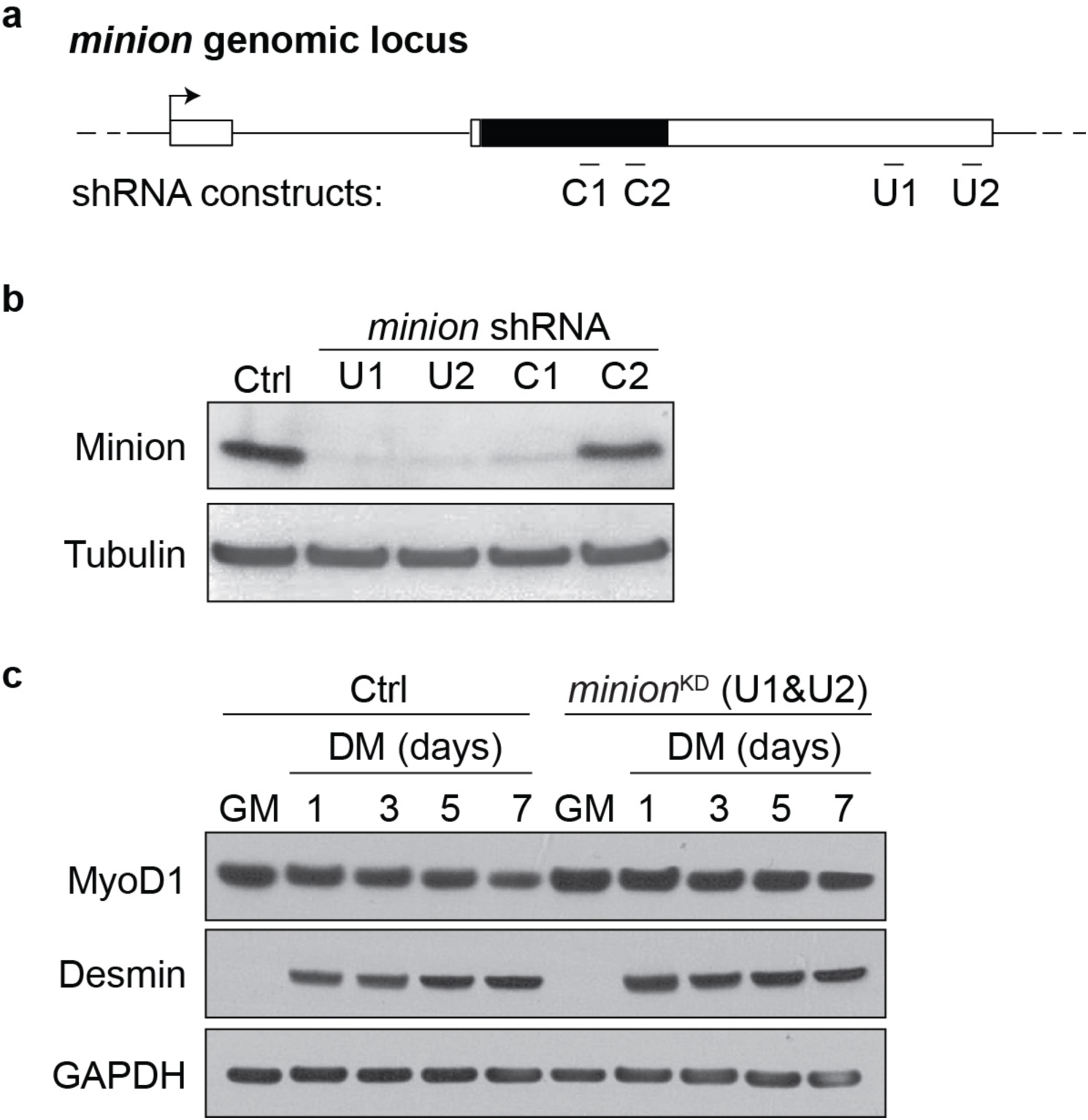
Generation and validation of lentiviral shRNA constructs targeting mouse *minion*. **(a)** Schematic of the mouse *minion* genomic locus with the target regions of four shRNA constructs underlined. Solid black bar indicated the *minion* ORF, and white bar indicated untranslated regions (UTRs). C1 and C2: shRNA constructs targeting the coding sequence; U1 and U2: shRNA constructs targeting the 3’UTR. **(b)** Western blot analysis of C2C12 myoblasts transduced with the indicated lentiviral shRNA constructs. A shRNA construct targeting the firefly (*Photinus pyralis*) *luciferase* gene was used as a negative control (Ctrl). After lentiviral infection and GFP sorting, the cells were expanded and kept in differentiation medium for 5 days. The two shRNA constructs targeting the *minion* 3’UTR (U1/U2) were found to reduce minion expression most efficiently and for subsequent experiments in C2C12 and primary myoblasts, cells were infected with U1 and U2 shRNA viruses and sorted by GFP after each round of infection to generate *minion*^KD^ cells. Similarly, C2C12 cells were infected with the control virus in two rounds and sorted twice by GFP to generate the control cells (Ctrl) below. n=2. **(c)** Western blot analysis of Ctrl and *minion*^KD^ C2C12 cells cultured in either growth medium (GM), or in differentiation medium for indicated number of days with indicated antibodies. n=3. Data are an expanded version of those shown in Fig. 3c.

**Extended Data Fig. 12.**
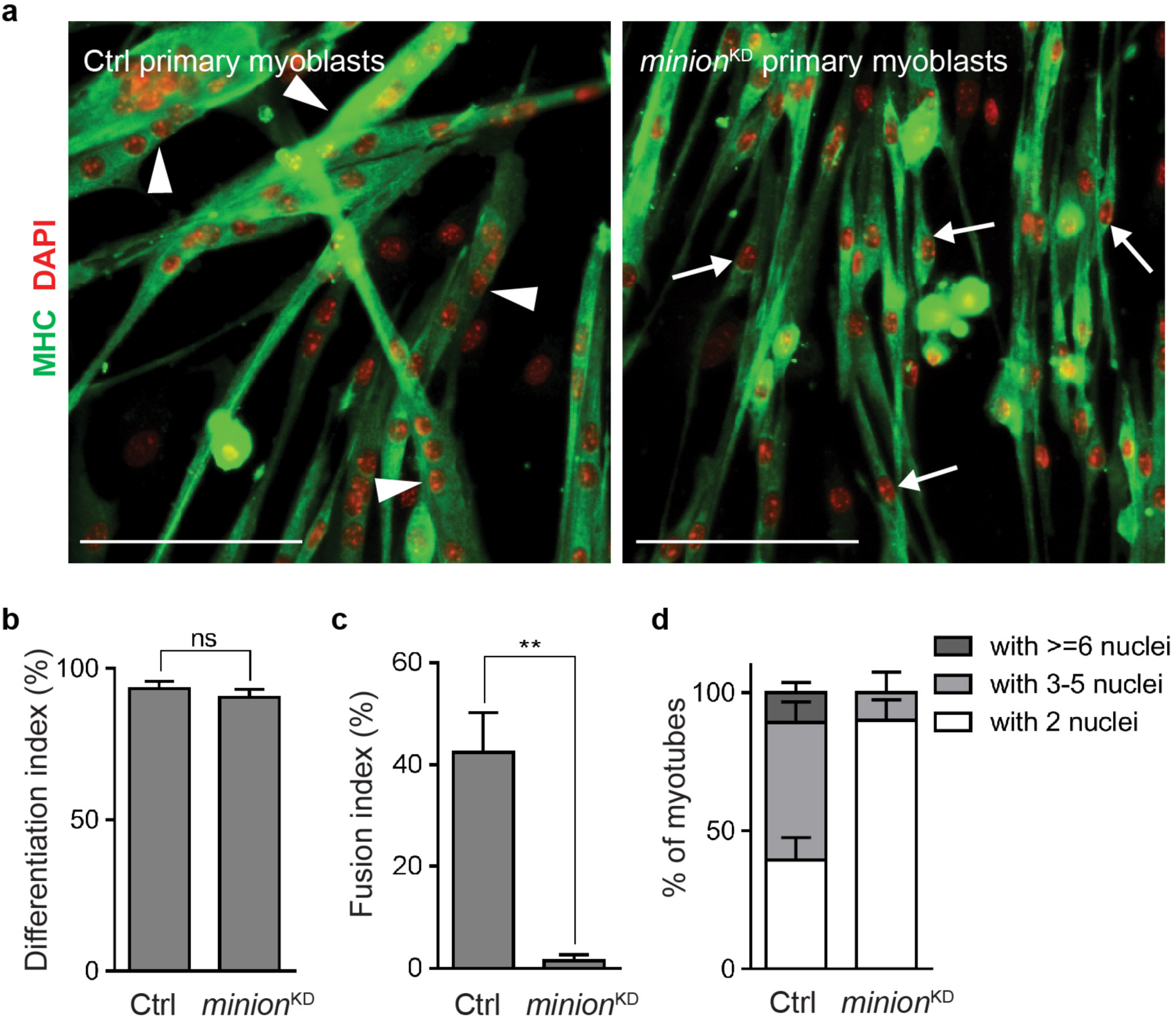
Knockdown of minion in primary adult myoblasts blocks myoblast fusion and polynucleated myotube formation. **(a)** Immunofluorescence images of Ctrl and *minion*^KD^ primary myoblast-derived myofibers, formed after 4 days in differentiation medium. White arrowheads and white arrows point out the fused multinuclear myotubes and unfused differentiating elongating myoblasts. n=2 (6 technical replicates each, 5 fields each replicate). MHC (green) and DAPI (red) staining are shown. Scale bar: 100 μm. **(b)** Quantification of differentiation index for the experiment done in (a). Differentiation index was calculated as the percentage of nuclei within MHC^+^ cells among total nuclei in each field. NS: not significant. **(c)** Quantification of fusion index for the experiment done in (a). Fusion index was calculated as the percentage of nuclei within MHC^+^ myotubes containing ≥3nuclei among total nuclei in each field **(d)** Quantification of the percentage of myotube numbers for the experiment done in (a). Myotubes were binned by nuclear number as indicated, and the percentage of myotubes within each subgroup was calculated. Double asterisks: *P* <0.001. b-d, n=6 (one 0.7mm×0.7mm field each).

**Extended Data Fig. 13.**
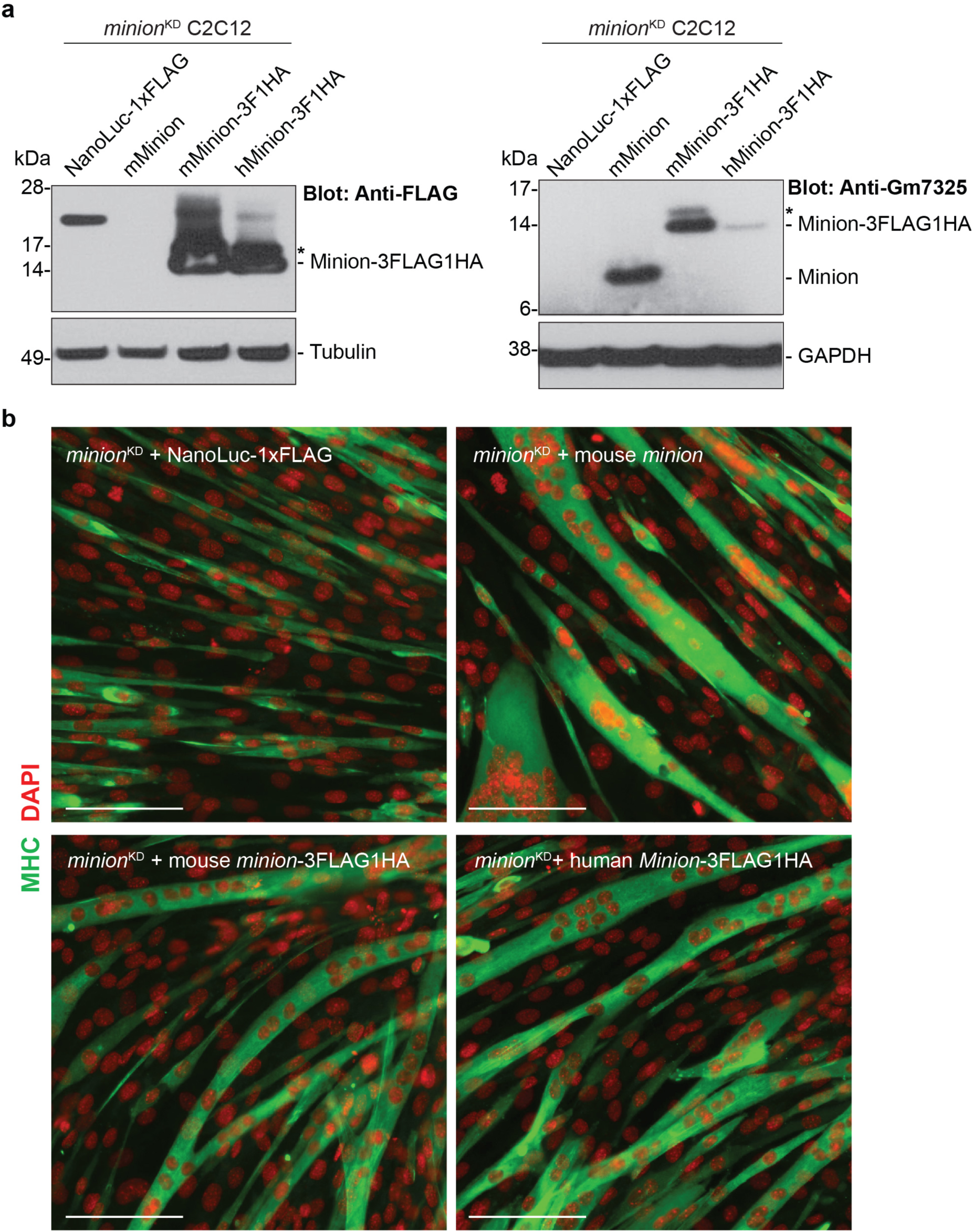
Exogenous expression of mouse minion or its human ortholog rescues the fusion defect of *minion*^KD^ myoblasts. **(a)** Western blots analysis of **minion*^KD^* cells after retroviral expression of untagged and C-terminally tagged minion and culture in differentiation medium for 5 days. Retroviral vectors carrying full-length mouse *minion* CDS, C-terminally 3×FLAG-1×HA-tagged mouse *minion* CDS and C-terminally 3×FLAG-1×HA-tagged human *Minion* ortholog CDS were used for reconstitution. A C-terminally 1×FLAG-tagged Nanoluc retroviral vector was used as negative control. The anti-FLAG antibody recognizes three tagged proteins at the correct size, while the anti-minion antibody not only recognizes the mouse minion protein (tagged and untagged) but also weakly recognizes the human Minion ortholog. Note: the pCIGAR gateway retroviral vectors carrying tagged human and mouse *Minion* CDS inherited an extra start codon at the 5’ end of and in-frame with the genuine start codon, giving rise to an extra band of slightly larger size (16 aa larger, asterisk). n=2. **(b)** Immunofluorescence images of *minion*^KD^ C2C12 cells with exogenous expression of tagged NanoLuc, untagged mouse minion, tagged mouse minion and tagged human Minion ortholog after 5 days in differentiation medium. MHC (green) and DAPI (red) staining are shown. n=2 (8 technical replicates each). Data are an expanded version of those shown in Fig. 3H. Scale bars: 100 μm.

**Extended Data Fig. 14.**
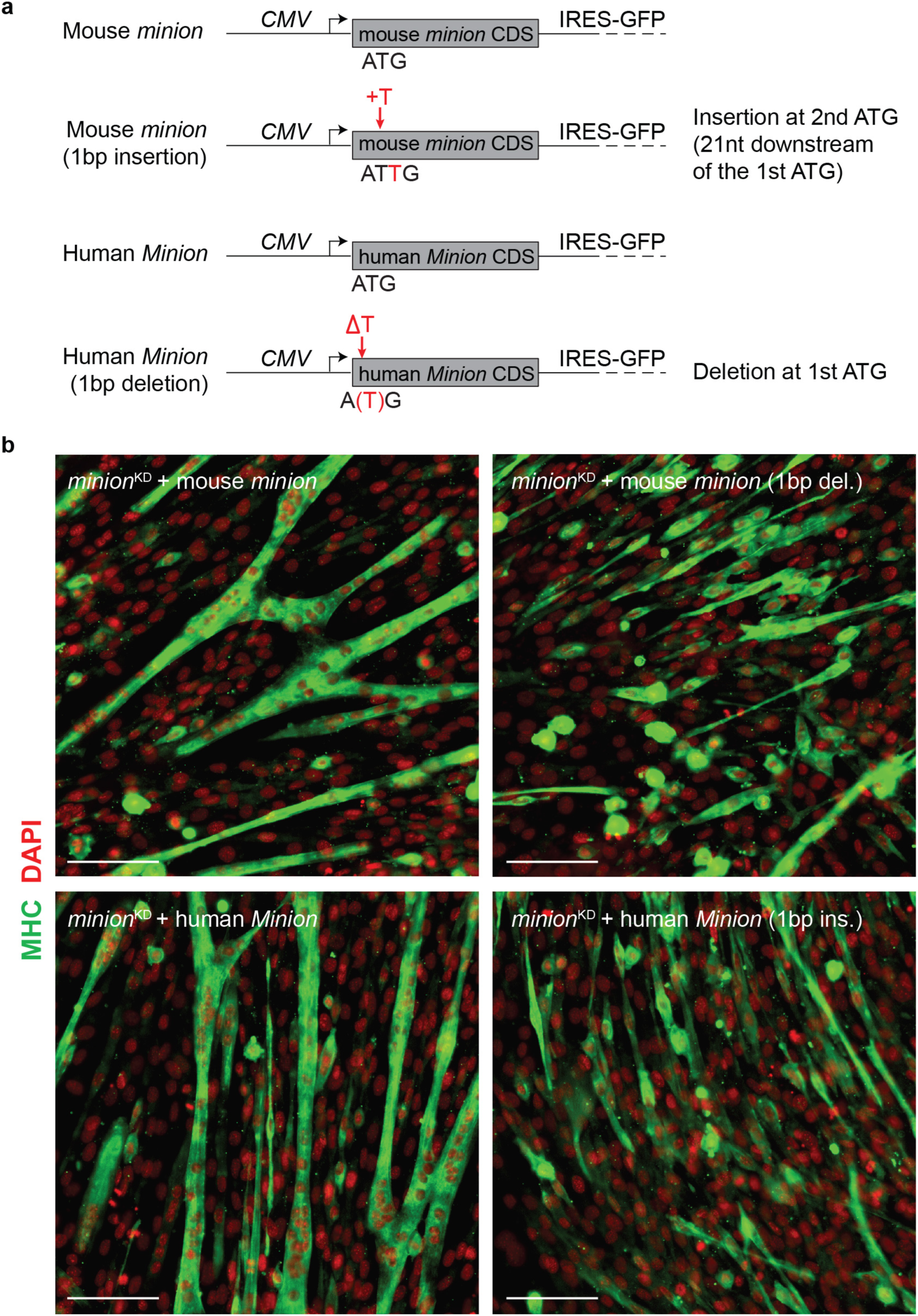
Mouse and human *minion* transcripts do not function as lncRNAs. **(a)** Schematic of four retroviral vectors containing either intact mouse/human *Minion* CDS, or those with 1bp frameshift (FS) mutations within the start codons (indicated in red). These single base pair mutations are predicted to disrupt the expression of full-length mouse/human Minion proteins without significantly altering RNA sequence. **(b)** Immunofluorescence images of *minion*^KD^ C2C12 cells with exogenous expression of constructs indicated in (a), and following 5 days in differentiation medium. MHC (green) and DAPI (red) staining are shown. n=2 (6 technical replicates each). Scale bars: 100 μm.

**Extended Data Fig. 15.**
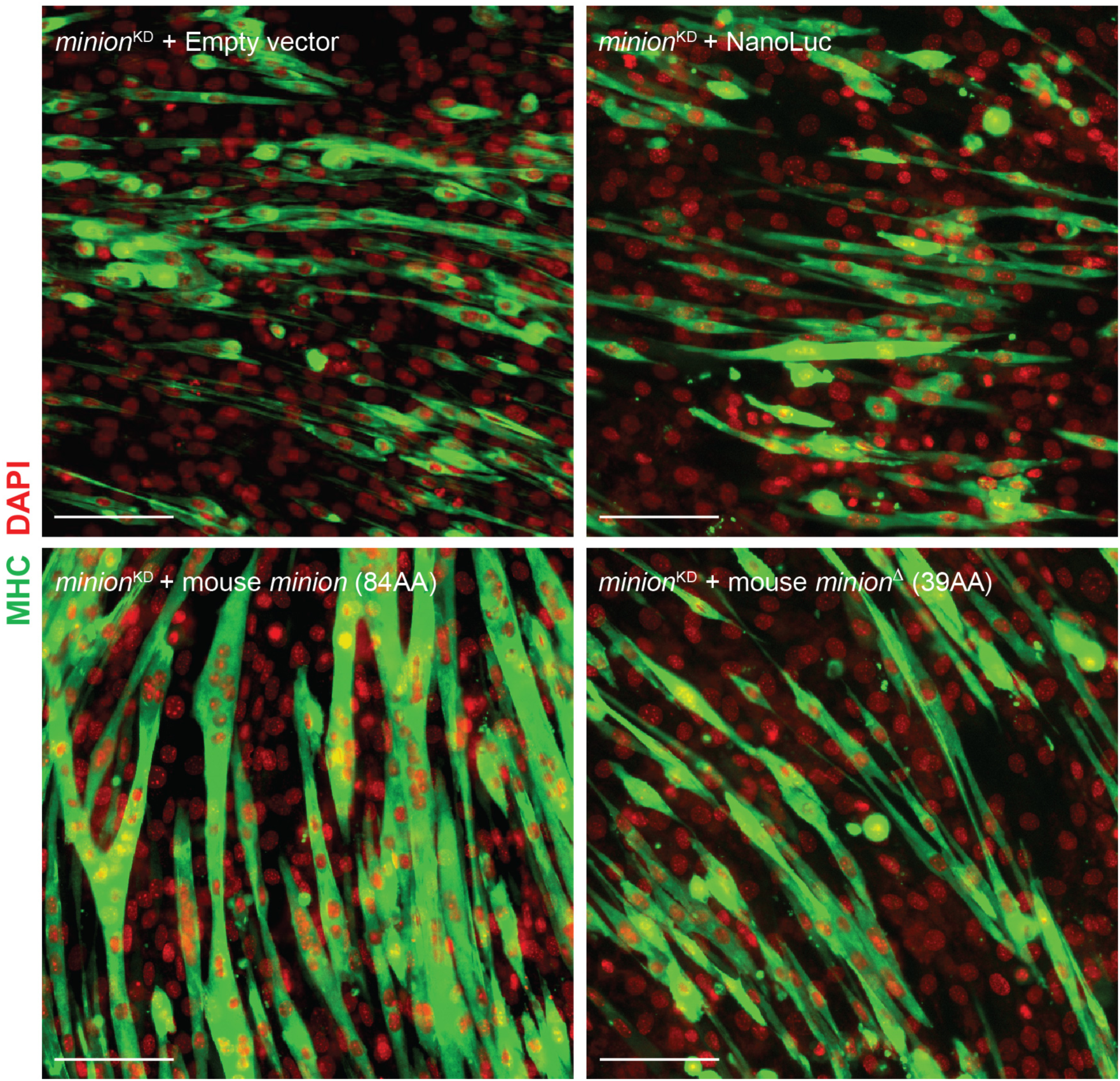
The minion^Δ^ truncation mutant predicted from the *minion* ^Δ^ knockout allele does not rescue the fusion defect observed in *minion*^KD^ C2C12 cells. Immunofluorescence images of *minion*^KD^ C2C12 cells transduced with retroviral constructs encoding either empty vector (negative control), NanoLuc (negative control), full-length mouse minion, or the truncated 39 aa minion mutant form (predicted from the *minion*^Δ^ knockout allele containing the 135bp in-frame deletion). Cells were cultured in differentiation medium for 5 days. MHC (green) and DAPI (red) staining are shown. n=2 (8 technical replicates each). Scale bars: 100 μm.

**Extended Data Fig. 16.**
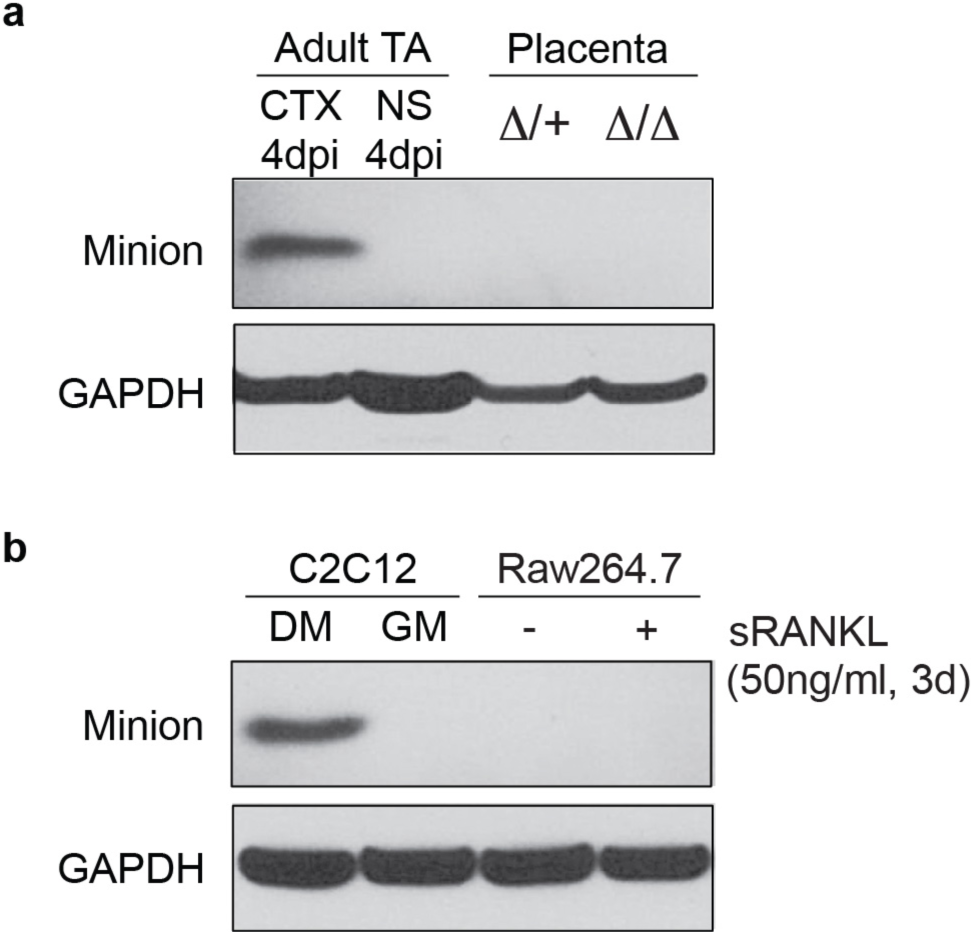
minion is undetectable in two models of non-myogenic cell-cell fusion. **(a)** Western blot analysis of minion expression in placenta. Placenta and periplacental tissue from *minion*^Δ/+^ and *minion* ^Δ/Δ^ embryos were examined. TA muscles with normal saline injection (NS; day 4) and cardiotoxin injection (CTX; day 4) were used as negative and positive controls, respectively. **(b)** Western blot analysis of minion expression upon soluble RANKL ligand-induced cell-cell fusion and osteoclast formation in the macrophage line Raw264.7. n=2. C2C12 cells cultured either in growth medium (GM), or in differentiation medium for 5 days (DM) were used as positive and negative controls respectively.

**Extended Data Fig. 17.**
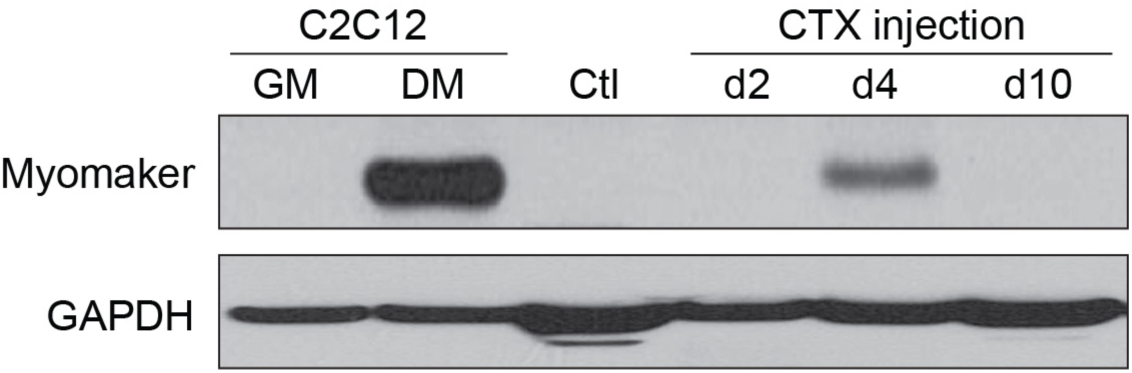
An anti-human Myomaker antibody recognizes endogenous mouse myomaker. Western blot analysis of C2C12 myoblasts cultured under either growth conditions (GM), or in differentiation conditions for 3 days (DM); and of uninjured (Ctl) or cardiotoxin injured and regenerating TA muscle at different time points (day 2, 4, 10). Antibody incubations were performed as described in Materials and Methods. n=2.

**Extended Data Fig. 18.**
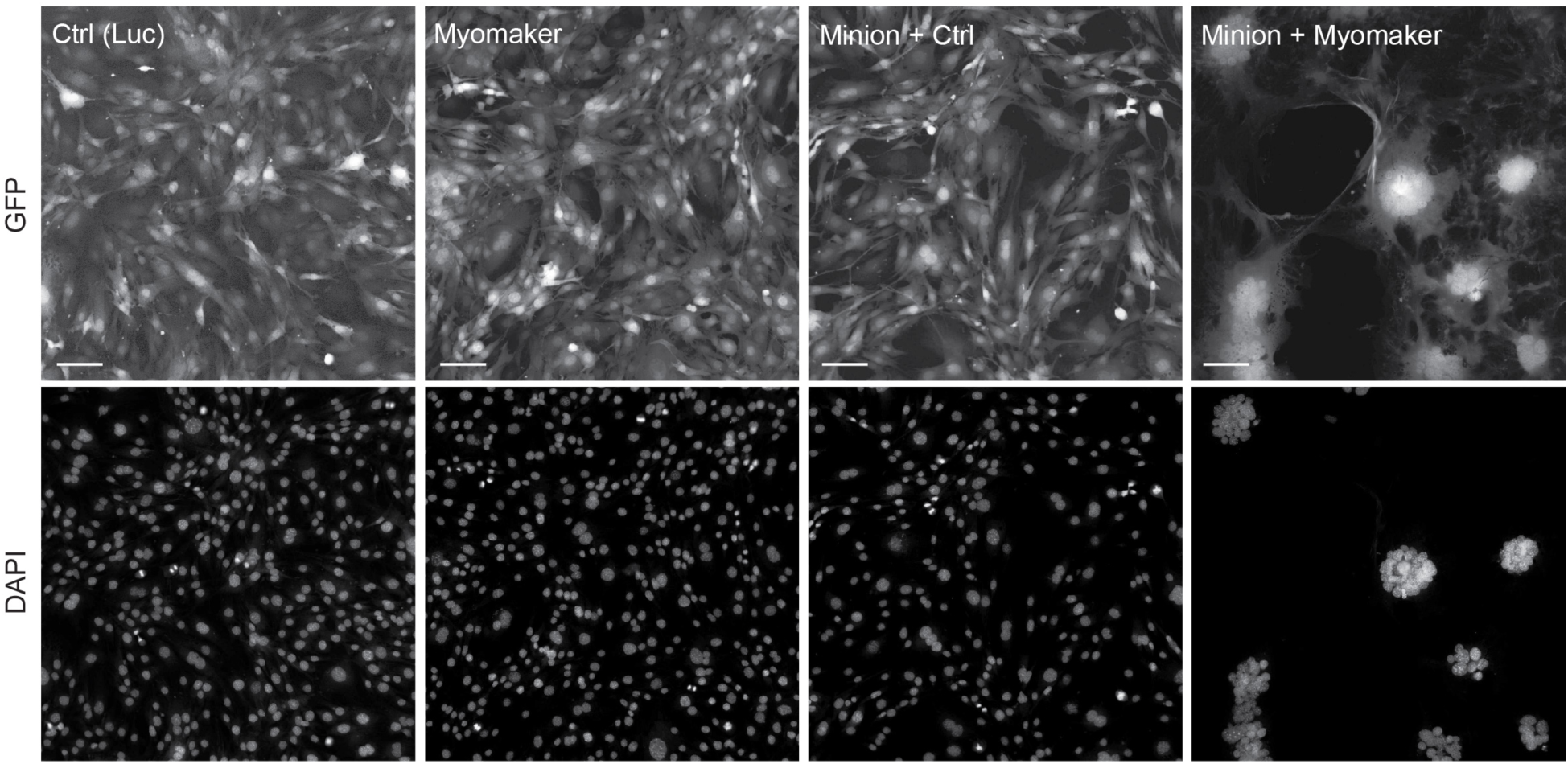
Minion and myomaker are together sufficient to induce multinuclear syncytia formation in 10T1/2 fibroblasts. Retroviral vectors encoding Luciferase, myomaker, or minion were transduced as indicated into 10T1/2 fibroblasts. All vectors contain IRES-GFP downstream of the gene of interest, causing infected cells to uniformly express GFP. Split-channel grayscale images for GFP and DNA are included. n=3 (8 technical replicates each). Fusion index was calculated as the percentage of nuclei found within GFP-positive syncytia containing ≥4 nuclei. Data are quantified in Fig. 4e. Scale bars: 100 μm.

**Extended Data Fig. 19.**
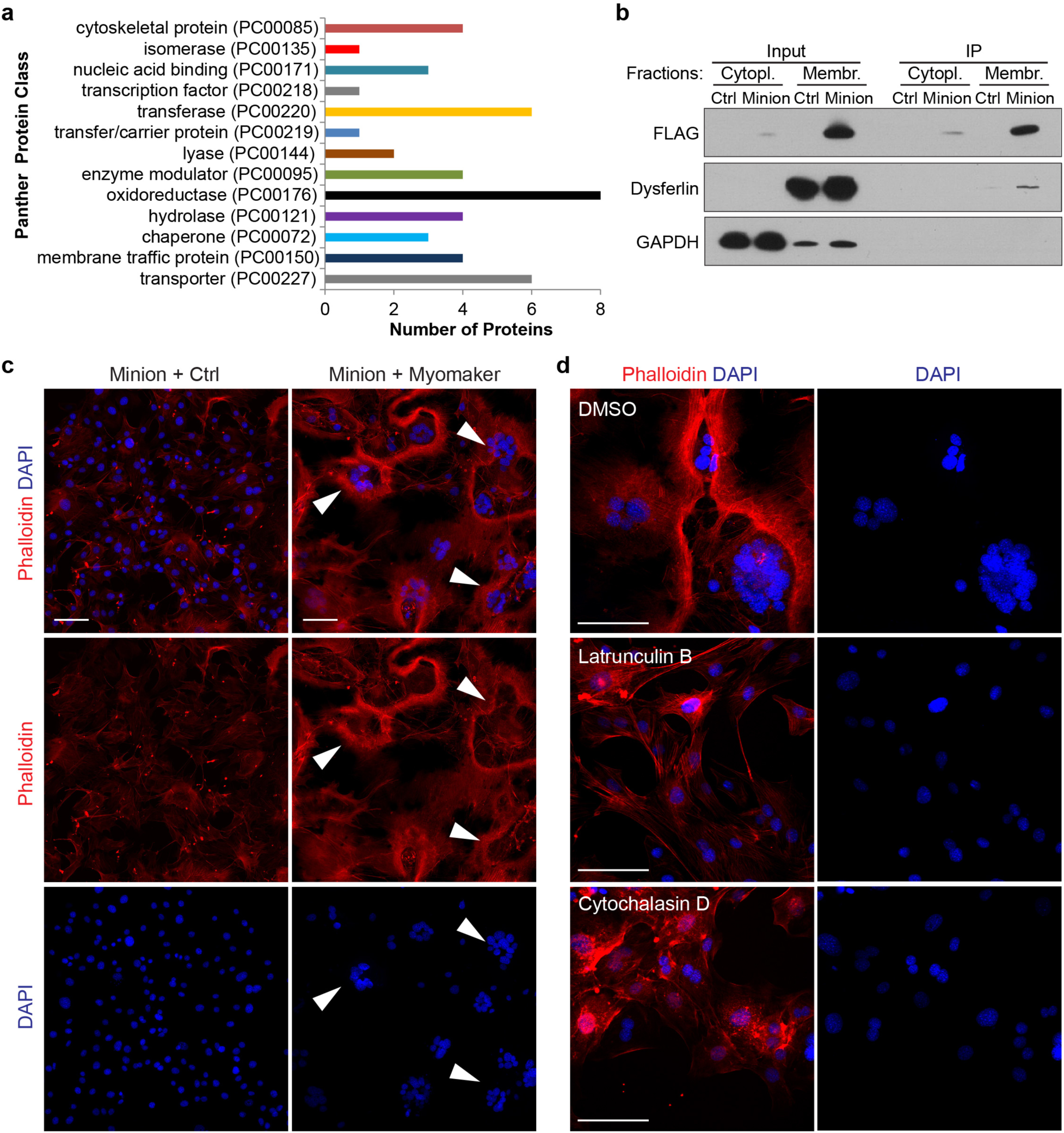
Fibroblast fusion induced by minion and myomaker requires actin cytoskeleton reorganization. **(a)** Minion-associated proteins were identified by AP-MS analysis from day 3.5 differentiating C2C12 myoblasts expressing FLAG-tagged minion (see Extended Data Table 1), and were grouped into protein classes using Panther.^32^ The number of significantly enriched proteins in each class is indicated. n=3. **(b)** Western blot confirmation of an example hit. Minion and GAPDH serve as positive and negative controls, respectively. **(c)** Fluorescence images of 10T1/2 fibroblasts co-overexpressing minion and Myomaker. F-actin (Alexa546-Phalloidin, red) and DAPI (blue) staining are shown. White arrowheads point to the boundaries of multinuclear cells. n=2 (6 technical replicates each, 5 fields each). **(d)** Fluorescence images of 10T1/2 fibroblasts co-overexpressing minion and Myomaker and treated for 24 hours with DMSO control or the actin polymerization inhibitors latrunculin B (0.1 μΜ) or cytochalasin D (0.3 μΜ)^17^. n=2 (6 technical replicates each, 5 fields each). Scale bars: 100 μm.

**Extended Data Fig. 20.**
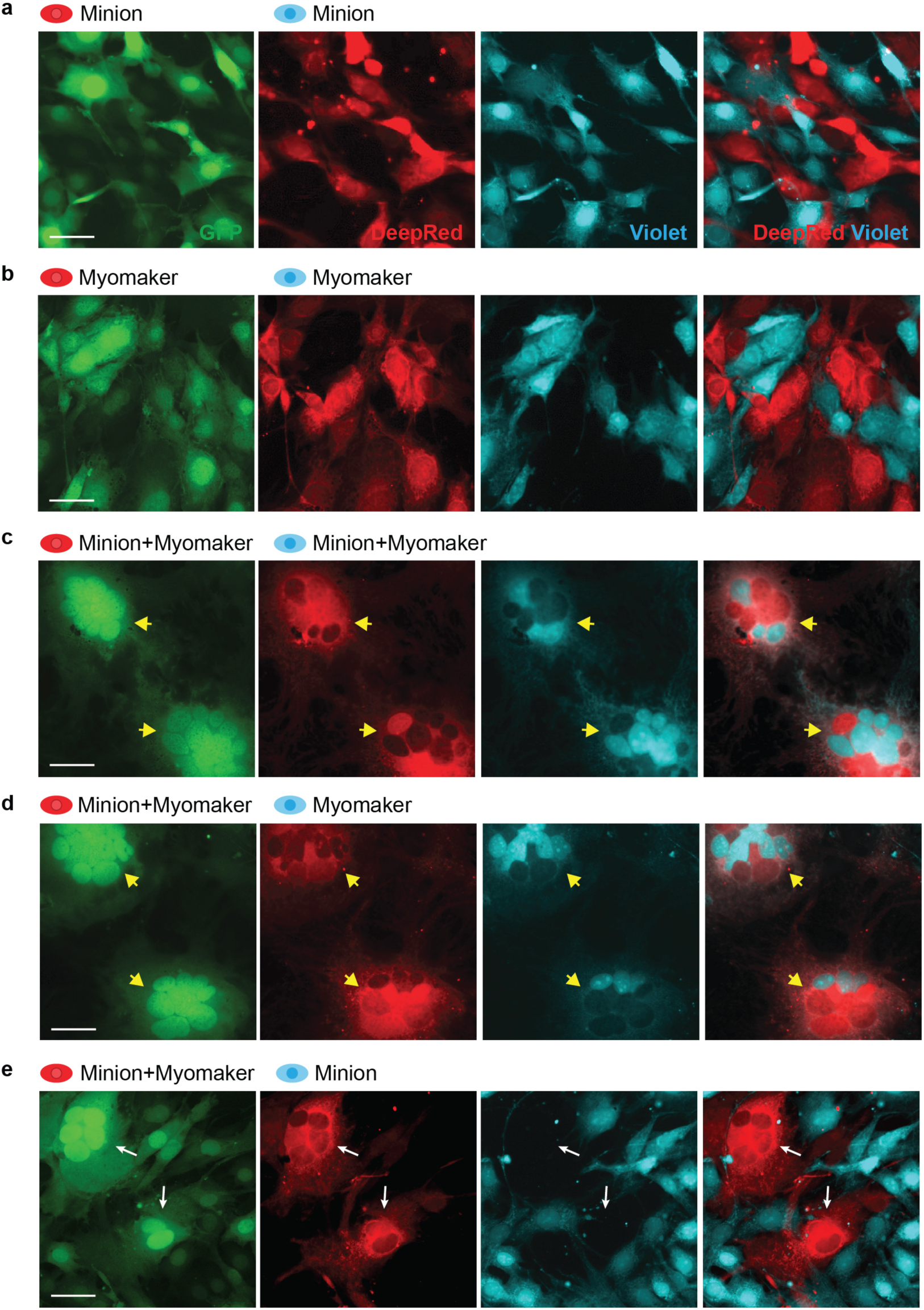
Minion and myomaker are together sufficient to induce cell-cell fusion in fibroblasts, with minion required only on one side of the fusion pair. Split-channel fluorescence images are provided of data included in Fig. 4f. Fluorescence images are shown from cell mixing experiments using fibroblasts expressing the indicated combinations of proteins and labeled with either CellTrace Violet (blue) and CellTracker Deep Red (red) dyes. 10T1/2 fibroblasts were serially infected with retroviruses encoding either minion, myomaker or control vectors (omitted in the labeling for simplicity). All vectors contain IRES-GFP downstream of the gene of interest, causing infected cells to express GFP. Relevant cell color and proteins expressed are indicated above each image. Yellow arrowheads indicate syncytia derived from Deep Red^+^ cells co-expressing minion and myomaker that also contain Violet^+^ nuclei from the second cell type expressing either myomaker only or minion and myomaker together. White arrows indicate syncytia derived from Deep Red^+^ cells co-expressing minion and myomaker that do not contain Violet^+^ nuclei from the second cell type expressing minion only. Scale bars: 50μm.

**Extended Data Table 1.**
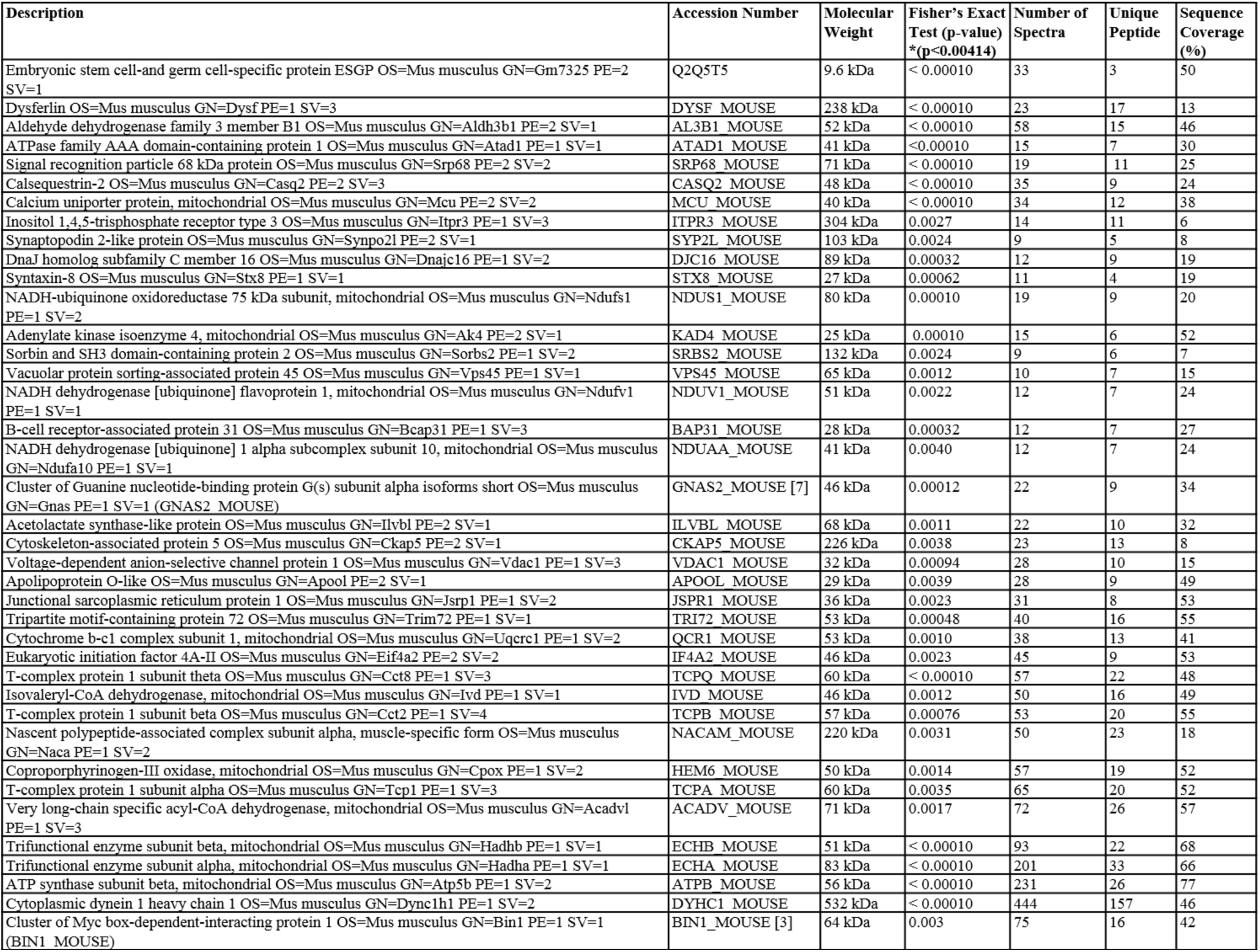
Proteins significantly enriched in complex with minion following affinity purification-mass spectrometry.

## Methods

### Animals

All animal experiments were approved by the GNF IACUC, and carried out in accordance with approved guidelines. C57BL/6J mice were initially purchased from the Jackson Laboratory and expanded through in-house breeding. For details regarding generation of genetically engineered mice, please see “Generation of *minion*-knockout mice by CRISPR/Cas9-mediated gene editing” below.

### Cardiotoxin (CTX) injury model

The cardiotoxin (CTX) injury model is a well-established model to study mouse skeletal muscle regeneration. CTX from *Naja mossambica mossambica* (Sigma C9759) was dissolved in normal saline (0.9% w/v of NaCl) to make a 10 μM working solution, and was aliquoted and stored at −20°C. After anesthesia of the mouse with isoflurane (1.5%-2% in oxygen), the anterior aspect of the adult mouse hindlimb (8- to 10-week-old C57BL/6) was sterilized with 70% ethanol, shaved to expose the skin, and approximately 50 μL of CTX solution was injected into the midbelly of the tibialis anterior (TA) muscle using a 0.3ml U100 BD insulin syringe. TA muscles were collected and examined at different time points after CTX injection. Adult mice at a similar age if not from the same litter without CTX injection or with equal volume normal saline injection were used as controls, as indicated in the figures.

### Mouse skeletal muscle RNA sequencing

Twelve 8- to 10-week-old C57BL/6 mice were injected with CTX into the TA muscle as described above, and the TA muscles were collected at 1, 3, 5, 7 days after CTX injection respectively with 3 mice for each time point. TA muscles from three 8- to 10-week-old uninjured mice were also collected. Total RNA from each muscle sample was isolated by TRIzol Reagent (Thermo Scientific 15596026) according to manufacturer’s instructions and purified by Qiagen RNeasy columns. The RNA samples (3 replicates for each time point) were submitted to the inhouse Sequencing and Expression Analysis Core for quality checking, library preparation, and next-generation single-read sequencing using standard techniques. Briefly, one microgram of total RNA was used to make Illumina-compatible sequencing libraries, and the libraries were sequenced using 50 bp single reads on an Illumina HiSeq 1000. Reads were aligned to the mouse transcriptome (Refseq mouse transcripts as of March 2013) using BWA^33^. An average of 36 million reads per sample mapped to the mouse transcriptome. To analyze the raw data, reads per kilobase of transcript per million mapped reads (RPKM) were calculated for each gene at each time point, the RPKM of each gene at CTX day 1, 3, 5, 7 was normalized to that of uninjured muscles, and the results were averaged to generate the fold change in expression level. The data were then analyzed in two ways: 1. Genes that exhibited more than 100-fold increase in CTX day 3 muscles compared to uninjured muscles were selected; 2. Genes annotated to contain an open reading frame (ORF) of less than 100 codons were selected. Genes meeting both criteria were then further examined against in-house RNA-Seq data from undifferentiated and differentiated primary myoblasts and C2C12 immortalized myoblasts; *minion* was the only small ORF demonstrating a dynamic expression pattern, with greater than 10-fold change between undifferentiated and differentiated myoblast samples.

### Developmental RNAseq analysis

RNAseq analysis of early embryonic development was from the Deciphering the Mechanisms of Developmental Disorders progam via EMBL-EBI Expression Atlas (https://www.ebi.ac.uk/gxa)^29,30^. The dataset was interrogated using the following Ensembl IDs: mouse *minion/gm7325*, ENSMUSG00000079471; mouse *myod1*, ENSMUSG00000009471).

### Generation of *minion*-knockout mice by CRISPR/Cas9-mediated gene editing

4-week-old female C57BL/6J mice were superovulated by intraperitoneal injection of 5 IU pregnant mare’s serum gonadotropin (PMSG) followed 47 hours later by 5 IU of human chorionic gonadotropin (HCG). Female mice were mated to C57BL/6J male mice 1:1 immediately after HCG injection. The following morning, the females were checked for copulatory plugs and zygotes were collected from the oviducts of plugged females. *In vitro* transcribed Cas9 mRNA (100 ng/ μL) and two gRNAs (50 ng/ μL) were coinjected into the pronuclei of fertilized zygotes. Zygotes surviving the injection procedure were transferred into a single oviduct of pseudopregnant ICR recipient females (50-60 embryos/oviduct). Mice produced from injected embryos were genotyped and sequenced (see “Assay for genome modification” below) to determine the presence of mutations within the genomic region of *minion*. Mutant founder animals were then bred to C57BL/6J mice and offspring were analyzed for germline transmission.

In order to generate *in vitro* transcribed Cas9 mRNA, a 10 bp spacer and the T7 promoter were added to the *Streptococcus pyogenes* Cas9 coding region by PCR amplification from a construct (pCR-Blunt II-TOPO-NLS-Cas9-NLS) made in-house, and the amplified gel-purified Cas9 PCR product was used as the template for *in vitro* transcription using mMESSAGE mMACHINE T7 ULTRA kit (Thermo Scientific).

The gRNA sequences were designed to target mouse *gm7325/minion* gene (Gene ID: 653016; Ensembl ID: ENSMUSG00000079471). In order to generate *in vitro* transcribed gRNA, two oligonucleotides were first synthesized (IDT):

Oligonucleotide 1: 5’ TTAATACGACTCACTATAG-(gRNA protospacer)-GTTTTAGAGCTAGAAATAGCAAGTTAAAATAAGGCTAGTCG;

Oligonucleotide 2: 5’ AAAAAGCACCGACTCGGTGCCACTTTTTCAAGTTGATAACGGACTAGCCTTATTTTAACTT After oligonucleotide annealing and PCR amplification, the T7 promoter with an additional “G” at the 5’ end (see oligonucleotide 1) was added to the gRNA. The amplified gel-purified gRNA PCR product was used as the template for *in vitro* transcription using a MEGAshortscript T7 kit (Thermo Scientific). Both the Cas9 mRNA and the gRNAs were purified using a MEGAclear kit (Thermo Scientific) and eluted into RNase-free water. The gRNA protospacer sequences targeting the mouse *minion* ORF region were as follows: sgRNA 1: 5’ GGACCGGGCCGTCGTGGAGG; sgRNA 2: 5’ CCAGAGTGGACCACTCCCAG.

### Assay for genome modification and genotyping

To detect mutations in the mice arising from injected embryos, PCR was performed using primers flanking the targeted region (Seq-F: 5’ GAGTGAACTCCTTAACCAGCTTTC, Seq-R: 5’ GCGTTGCTGTTTCCAGGACCCGTG). The PCR products were used in a surveyor assay according to manufacturer’s instructions (IDT). The PCR products were then analyzed by agarose gel electrophoresis and selected products were cloned into a pCR-blunt-cloning vector and sequenced. Mice containing mutations in the target region were bred to confirm germline transmission. Among the mouse strains with mutations, one with a 135 bp in-frame deletion within the *minion* ORF was selected for further analysis.

For genotyping of the subsequent progeny carrying the *minion*^Δ^ 135bp deletion allele, a primer set flanking the targeted region (F0: 5’-CAAAGGGAGGGAGGGATTAAAG-3’; R0: 5’-CAGAGAGGAAG GGTCAATCAAC-3’) was used to amplify genomic DNA, generating a ∼760 bp product from the unmodified allele and a ∼625 bp product from the mutated allele respectively. The PCR products were separated by gel electrophoresis using 2% agarose (Sigma A9539). Wild-type mice demonstrate a single band of the larger size, while homozygotes containing the deletion demonstrate a single band of the smaller size, and heterozygous mice demonstrate both bands.

### Myoblast isolation

For mouse embryonic myoblast isolation, pregnant C57BL/6J female mice were humanely euthanized, and embryos were rapidly but gently dissected and placed into dissection buffer containing Ham’s F10 nutrient mix (Thermo Scientific 11550043) with 1x antibiotic-antimycotic (Thermo Scientific 15240062). For each embryo, the tail was kept in a numbered tube for genotyping, while all four limbs were skinned, dissected, and placed into a 2ml numbered Eppendorf tube containing ∼1.5 ml dissection buffer including 4 mg/ml Collagenase type II (Worthington LS004176; freshly made and filtered before use). The samples were rotated on a platform rocker at 80 rpm and 37 °C for 30-45 min, until the muscles were mostly digested and only bones and soft tissues were left. In general, earlier stage embryos required shorter incubation times. Cell suspensions were checked microscopically after each step to avoid over-digestion. After allowing the unwanted tissues to settle at room temperature, the supernatant was transferred to a 50ml conical tube. Dispase II (Thermo Scientific 17105041) was then added at a final concentration of >=1.2 mg/ml (>0.6 U/ml) to both tubes: (1) For the supernatant in the 50ml tubes, cells were incubated at 37 °C for 20-30 min with occasional mixing; (2) For the remaining tissues in the 2ml tubes, 1.5ml freshly made and filtered dissection buffer with 4 mg/ml Collagenase type II and >=1.2 mg/ml Dispase II, and the tubes were again rocked on the rocker at 80 rpm and 37 °C for 20-30 min to allow further digestion and dissociation.

After dissociation, the suspension in the 2ml tube was mixed with that in the 50ml tube. After adding ∼4-5 volumes of wash medium (Ham’s F10 and 10% horse serum, filtered), the digested mixture was passed through a 10 ml 20 gauge needle slowly and gently for approximately 4 times, while scrupulously avoiding generation of bubbles. More wash medium was then added to bring the final volume to 30ml. This suspension was filtered through a prewashed 40 μm Nylon Mesh filter on top of a new 50ml conical tube, and the filter was rinsed with 10 ml wash medium into the same tube. All of the 50 ml tubes were then centrifuged at 125 × g for 5 min at room temperature, and the supernatant was transferred and spun down again at 125 × g for 5 min. The pellets from two centrifugations were resuspended and mixed in 2 ml myoblast isolation medium followed by an additional 20 ml of media containing a 1:1 mixture of DMEM low glucose (Gibco 11885084) and Ham’s F-10 Nutrient Mix (Gibco 11550043); 20% (v/v) FBS; 1x antibiotic-antimycotic; and freshly added 2.5 ng/ml rhFGF (Promega G5071). Medium lacking DMEM but containing the remaining items above also produced similar results.

The isolated cell mixture from each embryo was first plated into a regular 150 mm TC-treated dish for 30 min at 37 °C (preplate I) and replated into another 150 mm dish for 30min at 37 °C (preplate II) in order to eliminate fibroblasts, and then the supernatant containing mostly myoblasts was transferred into two 100 mm collagen-coated dishes. Cells were examined the next day to determine necessity for passaging. Occasionally the preplate II dish also contained some amount of myoblasts, and these were kept and expanded in addition to those in the collagen dishes. 0.05% trypsin was used for dissociating the cells from dishes. After a few passages, 1x antibiotic-antimycotic was replaced by 1x penicillin-streptomycin (Gibco). As the fibroblast number decreases in culture, the embryonic myoblasts may start to proliferate very slowly and they should be seeded more densely to recover from the slow growth. Adult mouse myoblasts were isolated similarly but with a few modifications: the muscles were removed from bones and minced with small scissors; 15 ml conical tubes were used instead of 2 ml tubes, with twice the volume of digestion buffer; and longer digestion and dissociation times were used.

### *In vitro* myoblast differentiation assay

For primary myoblasts derived from both embryos and adult mice, approximately 3000 cells in 50 μl myoblast growth medium (1:1 mixture of DMEM low glucose and Ham’s F-10 Nutrient Mix; 20% (v/v) FBS; freshly added 2.5 ng/ml rhFGF) were seeded into each well of a 384-well Collagen-coated PerkinElmer CellCarrier plate (6007550) for imaging purposes. The next day, differentiation medium (DMEM high glucose (Gibco 11995073) with 3% to 5% horse serum) was added to the cells (DM day 0). Differentiation medium was replaced daily. The cells were fixed at DM day 3 and day 4 for immunofluorescence staining. For C2C12 cells, ∼ 1500-2000 cells were seeded into each well of a 384-well plate (DMEM high glucose with 10%FBS) and around 2 × 10^5^ cells were seeded into each well of a 6-well plate, using C2C12 growth medium (DMEM high glucose with 10%FBS). The following day, differentiation medium (DMEM high glucose with 2% horse serum) was added to the cells (DM day 0), and differentiation medium was subsequently replaced daily. The cells were collected or fixed at different time points as described.

### Histology

For paraffin sections with embryonic samples, mouse embryos (E14.5 and later) were decapitated and the tails were collected in numbered tubes for genotyping. To enhance fixation in later-stage embryos (E17.5 and later), embryos were skinned in the area to be studied. Embryos were fixed overnight using 4% paraformaldehyde (PFA; Electron Microscopy Sciences #15714) in PBS at 4 °C with gentle rotation. Following two quick rinses with PBS, embryos were placed into 70% ethanol for dehydration and long-term storage. For the tissues/organs to be studied, the appropriate portions were cut and submitted to histology core for paraffin embedding and sectioning using routine protocols.

For cryosections of adult mouse tissue, skeletal muscle samples were dissected and partially embedded in gum tragacanth (Sigma G1128; 10% w/v in PBS) on a wooden dowel, and frozen in 2-methylbutane in a glass beaker cooled on liquid nitrogen. The fresh frozen muscle samples were then sectioned at 10 μm thickness using a cryostat cooled to −20°C. These fresh frozen muscle sections were then fixed in 1% PFA diluted in PBS at room temperature for 5 min before subsequent staining procedures. Both cryosections and paraffin sections were stained with Hematoxylin and Eosin (H&E) following routine protocols.

### Immunofluorescence staining on tissue sections

For muscle cryosections, after fixation with 1% PFA/PBS as described above, slides were washed with PBS and permeabilized with 0.2% Triton X-100 diluted in PBS at room temperature for 10 min, and were then washed again with PBS. Sections were blocked at room temperature for 1 hour using a freshly-prepared and filtered solution containing 1% heat-inactivated donkey serum, 1% BSA, 0.025% Tween20 in PBS. After blocking, sections were incubated with primary antibody at 4 °C overnight, washed with PBS, and then incubated with secondary antibody for 2 hours at room temperature. After a 5 min wash with PBS, the sections were incubated with the nuclear stain DAPI (Molecular Probes D1306; 5 mg/ml stock) at a 1:20,000 dilution in PBS for 5 min, and slides were mounted and sealed using ImmuMount (Shandon) and glass coverslips. For paraffin sections with embryonic tissues, the deparaffinized and rehydrated slides were permeabilized with 0.2% Triton X-100 in PBS for 10 min and washed again with PBS. The sections were then blocked at room temperature for 1 hour using a freshly-made and filtered solution containing 5% heat inactivated normal goat serum in PBS. After blocking, similar procedures were performed as mentioned above for cryosections.

Primary antibodies used for immunofluorescence were: Mouse anti-MHC (MY32 clone, Sigma M4276, 1:300 dilution on paraffin sections and 1:500 dilution on cryosections); Mouse anti-Desmin (D33 clone, DAKO M0760, 1:300 dilution); Sheep anti-Gm7325/minion (R&D systems AF4580; 1:200 dilution on cryosections only). All secondary antibodies (Invitrogen Alexa-Fluor) were used at 1:250 dilution, and the host species was either donkey or goat. Only secondary antibodies from the same host species were used together for co-staining.

### Immunofluorescence staining and fluorescence staining with adherent cells

For immunofluorescence staining with adherent cells, mainly C2C12 and primary myoblasts from adults and embryos, 384-well PerkinElmer CellCarrier plates were again used. Cells were fixed with 4% PFA in PBS for 8-10 min and quickly washed twice with PBS before permeabilization with 0.2% Triton X-100 in PBS for 10 min. After one wash with PBS, cells were blocked with freshly made and filtered 5% heat inactivated normal goat serum in PBS for 1 hour, and were incubated with primary antibodies overnight at 4°C. The next day, after two quick washes with PBS, cells were incubated with secondary antibodies for 1-2 hours at room temperature. After three quick washes with PBS, the cells were incubated with DAPI (5 mg/ml stock, 1:20,000 dilution in PBS) for 10 min. The 384-well plate was then imaged using either UltraVIEW confocal or ImageXpress Micro (IXM; Molecular Devices) confocal imaging systems (see the Microscopy part below).

Primary antibodies used were: Mouse anti-MHC (MY32 clone, Sigma M4276, 1:400 dilution); Mouse anti-Desmin (D33 clone, DAKO M0760, 1:300 dilution). All secondary antibodies (Invitrogen Alexa-Fluor) were used at 1:250 dilutions, and the host species was either donkey or goat. Only secondary antibodies from the same host species were used together for co-staining.

For fluorescence staining of actin filaments in fibroblasts and myoblasts, the high-affinity F-actin probe Alexa Fluor 546-conjugated phalloidin (Invitrogen A22283) was used according to manufacturer’s instructions. Briefly, cells grown in 384-well plates were fixed with 4% PFA for 10 min at room temperature, washed with PBS, and permeabilized with 0.1% Triton X-100 for 5 min. After two PBS washes, the cells were blocked with PBS containing 1% BSA for 30 min and incubated with staining solution (1:80 dilution of the phalloidin methanolic stock in blocking solution) for 1 hour at room temperature. After two to three quick PBS washes, the cells were incubated with DAPI (5 mg/ml stock, 1:20,000 dilution in PBS) for 10 min. The plate was then imaged using either UltraVIEW confocal or IXM confocal imaging systems (see the Microscopy part below).

### Microscopy and imaging

The Invitrogen EVOS FL Auto Imaging System was used for routine examination of immunofluorescence staining, GFP virus infection, and cell labeling. For imaging of histological and immunostained tissue sections on glass slides, the Hamamatsu NanoZoomer and Aperio VERSA scanners were used to obtain whole-slide images using a 20x objective. For imaging of the immunofluorescence cell samples in 384-well plates, the IXM confocal high-content imaging system was used with 10x and 20x objectives. In order to acquire higher resolution images for tissue sections and cell samples, the UltraVIEW VoX 3D live cell imaging system (PerkinElmer) spinning disk confocal microscope system was used with 20x, 40x and 60x objectives. All pictures of whole mouse embryos were taken using iPhone 5S in combination with Leica KL200 LED dissection microscope.

### Cell culture

For culture of primary myoblasts isolated from later-stage mouse embryos and adult mice, filtered myoblast growth medium (1:1 mixture of DMEM low glucose and Ham’s F-10 Nutrient Mix; 20% FBS) with freshly added 2.5 ng/ml rhFGF was used. In general, around 2-4 × 10^5^ cells were seeded into a 100 mm collagen-coated dish, and the cells were split once every 2 to 3 days at a ratio of 1:2 to 1:4, depending on proliferation speed. 0.05% trypsin was used for dissociating cells from dishes. Myoblasts typically went through a crisis period after the removal of most fibroblasts, and could be seeded more densely at this point. Primary myoblasts in culture were monitored every day with fresh medium replacement as needed. For the culture of immortalized C3H/C2C12 myoblast cells (ATCC), filtered C2C12 growth medium (DMEM high glucose with 10%FBS) was used. Approximately 1.5 × 10^5^ cells were seeded into a 100 mm tissue culture-treated dish, and cells were split every 2 days. 0.25% trypsin was used for cell dissociation. The cells tested negative for mycoplasma contamination. For culture of the immortalized C3H/10T1/2 fibroblasts (ATCC), filtered fibroblast growth medium (DMEM high glucose with 15% FBS) was used. Approximately 1 × 10^5^ cells were seeded into a 100 mm tissue culture-treated dish, and the cells were split once every 3 days. 0.25% trypsin was used for cell dissociation. The cells tested negative for mycoplasma contamination. For culture of the immortalized RAW264.7 macrophage line (ATCC), filtered growth medium containing DMEM high glucose with 10% FBS was used. Around 2-3 × 10^6^ cells were seeded into a 175 cm^2^ flask. The cells were split once every 2 to 3 days, when they were ∼60-75% confluent. To ensure cell lifting and reduce cell death, 0.25% trypsin and a cell scraper were used in combination. To induce the formation of multinuclear osteoclast-like cells, 50 ng/ml sRANKL (Peprotech, 174aa) was incubated with the cells for 3 days. For the culture of CJ7 embryonic stem cells derived from 129 mice, freshly made and filtered growth medium was used, consisting of ESGRO Complete PLUS medium (Millipore SF001-500P) with 15% FBS and 3 inhibitors: GSK3β inhibitor which comes with the medium; MEK inhibitor PD184352 (0.8 μM final); and FGFR inhibitor PD173074 (0.1 μM final). Normally the cells were co-cultured with mouse embryonic fibroblasts according to standard procedures, but for the purpose of RNA and protein isolation they were seeded onto gelatin-coated dishes without a fibroblast feeder layer. Approximately 1 × 10^6^ cells were seeded into each 100mm dish. The medium was replaced every day. Cells were split once every two days at a ratio of 1:5 to 1:10, depending on experimental need. 0.05% trypsin was used for cell dissociation. All cell culture media contained 100 units/ml of penicillin and 100 μg/ml of streptomycin, unless otherwise specified. None of the cell lines is listed in the database of commonly misidentified cell lines maintained by ICLAC.

For experiments shown in Extended Fig. 19d, 10T1/2 cells of indicated genotypes were seeded into 384-well Perkin Elmer COC plates at 800, 1600, 3200 cells/well in fibroblast growth medium. After 15 hours, the cells were incubated with growth media containing DMSO (0.003%), latrunculin B (100 nM) or cytochalasin D (300 nM) for 24 hours before further analysis^17^.

### Tissue and cell lysates preparation for protein analysis

Both embryonic and adult mouse tissue samples were weighed, snap-frozen in liquid nitrogen, and stored at -80 °C until use. For preparation of protein lysates, eight volumes of ice-cold lysis buffer (50 mM Tris-HCl pH7.5, 150 mM NaCl, 1 mM EDTA, 10% glycerol, with freshly added 2x Halt protease inhibitor cocktail and 1x Roche PhosSTOP phosphatase inhibitor cocktail) and one to two 3 mm tungsten carbide beads (Qiagen) were added to each sample in a 1.5 ml or 2 ml Eppendorf tube. These were then homogenized at 30 cycles/s for 3-8 min at 4 °C using a TissueLyser II. Detergents were then added to the lysates to a final concentration of 0.1% SDS, 0.1% sodium deoxycholate and 1% Triton X-100, and the samples were rotated at 4 °C for 2-4 hours. Lysates were then transferred to new tubes and spun down at 1500021000 χ g for 10 min at 4°C. For organs containing significant amount of lipids, the supernatant was transferred and spun down again at 15000-21000 × g for 10 min at 4°C.

For cell samples, buffer from Alfa Aesar (J60423) was generally used (50 mM Tris-HCl pH7.5, 150 mM NaCl, 5%Glycerol, 0.1% SDS, 0.5% sodium deoxycholate and 1% Triton X-100, with the above mentioned protease and phosphatase inhibitor cocktails). Cells were quickly rinsed with DPBS and then ∼300 μL ice-cold lysis buffer was added to each well of a 6-well plate. After incubation on ice for 5 min, the cells were pipetted up and down and transferred to 1.5 ml Eppendorf tubes and incubated on ice for 30 min, with 1 sec of vortexing every 10 min. The samples were then spun down at 15000-21000 × g for 10 min at 4°C. A second buffer without ionic detergents was used for some of experiments (50 mM Tris-HCl pH7.5, 150 mM NaCl, 10% glycerol, 1 mM EDTA, and 1% Triton X-100, with the above mentioned protease and phosphatase inhibitor cocktails), with similar results. Bio-Rad DC protein assay with BSA standard was performed to measure protein concentration of supernatants. Lysates were mixed with NuPAGE LDS Sample Buffer (NP0007) and dithiothreitol (100 mM final), and boiled at 94°C for 10 min prior to SDS-PAGE.

### Subcellular fractionation analysis

C2C12 cells incubated with differentiation medium for 4 days were used for the subcellular fractionation studies using the Qproteome Cell Compartment system (Qiagen 37502) following the manufacturer’s instructions. The cells were dissociated from the dishes first before addition of the first buffer; similar results were obtained with direct cell lysis on the plate. The cytosolic/membrane/nuclear/cytoskeletal fractions were extracted from the cells, and for each fraction, the protein lysate extracted from an equivalent number of cells was loaded for Western blot (Extended Data Fig. 2d).

In order to examine whether minion protein is secreted and soluble in cell conditioned media, C2C12 myoblasts with indicated genotypes were first differentiated for two days in differentiation medium containing 2% horse serum, and then incubated with serum-free differentiation medium consisting of DMEM with 1x ITS-G (Thermo Scientific 41400045) for 24 hours. The supernatant conditioned media were collected, centrifuged at 8°C in order to eliminate dead cells, and filtered using a 0.45 μm vacuum filter bottle to further eliminate cell debris. The filtered supernatants were then concentrated at 8°C using 3k Amicon ultracentrifugation filters for multiple rounds according to the manufacturer’s instructions. The supernatants were concentrated approximately 400-1000 fold. As a comparison, whole cell extracts were prepared from the original cell pellet using the lysis buffer mentioned above, and denatured supernatants and whole cell extracts generated from equal amounts of cells were loaded for western blot analysis (Extended Data Fig. 2e).

### SDS-PAGE and Western blots

The NuPAGE Novex gel electrophoresis system was used for the separation of proteins. Approximately 10-30 μg of cell lysate or 30-60 μg of tissue lysate were loaded per well. NuPAGE MES SDS Running Buffer (NP0002) and 4-12% NuPAGE Novex Bis-Tris gels were used. Proteins were transferred to PVDF or nitrocellulose membranes using the iBlot transfer system (Thermo Scientific). Freshly prepared 5% milk in TBST (137 mM NaCl, 20 mM Tris, 0.1% Tween-20, pH7.6) was generally used as the blocking buffer with both PVDF and nitrocellulose membranes. However, for the detection of minion protein using primary antibody raised in sheep, freshly made and filtered 10% donkey serum in TBST was used as the blocking buffer with PVDF membrane (Millipore Immobilon-P^SQ^, 0.2 μm pore size). The information of primary and secondary antibodies used in Western blots is listed in the table below (Table 1). Two ECL substrates with different sensitivity were used as indicated. We found that the antihuman TMEM8C antibody recognized both endogenous and overexpressed myomaker protein in both mouse primary muscle and cultured cell lysates (Fig. 4c, d, Extended Data Fig. 17), but required extended antibody incubation and exposure times.

**Table 1.**
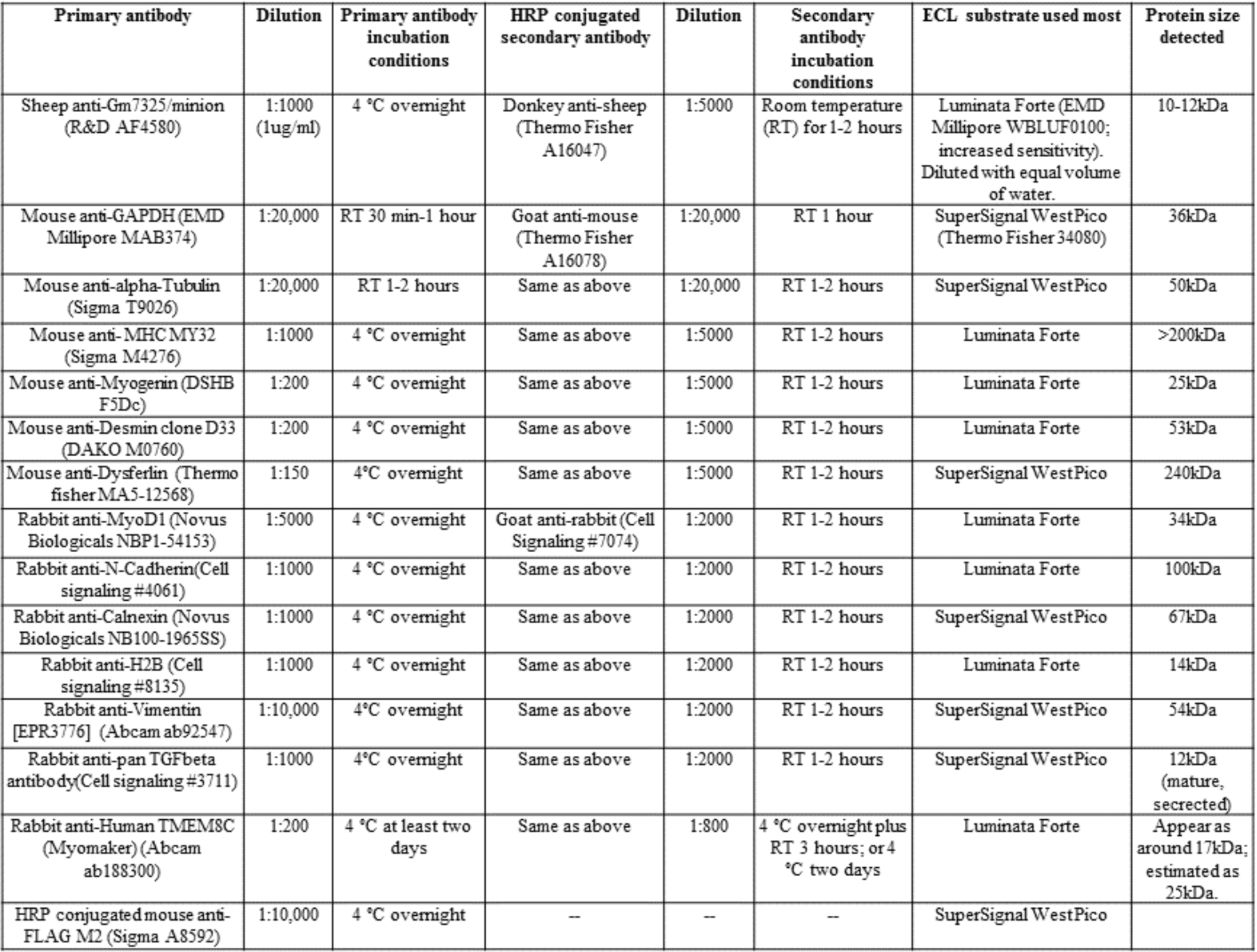
Antibody and ECL substrate information for Western blots.

### RNA preparation and RT-qPCR

Total RNA was isolated from cell lines using TRIzol Reagent (Thermo Scientific 15596026) according to the manufacturer’s instructions. First-strand cDNA synthesis was performed using qScript cDNA SuperMix (Quanta BioSciences) according to manufacturer’s instructions. For PCR, cDNA from ∼5 ng RNA was used in a 12.5 μl reaction with Power SYBR Green PCR Master Mix (Thermo Scientific 4367659). Reactions with RNA only were prepared as negative controls. An Applied Biosystems 7900HT thermocycler was used with the following primers: *minion* (5’-GGACCACTCCCAGAGGAAGGA-3’ and 5’-GGACCGACGCCTGGACTAAC-3’) and *gapdh* (5’-AGGTCGGTGTGAACGGATTTG-3’ and 5’-TGTAGACCATGTAGTTGAGGT-3’). Relative quantification was performed using the comparative CT method. The CT value of *minion* gene was normalized to that of the reference gene *gapdh* in the same sample using the formula: 2^ΔΔCT^.

### Lung flotation assay

Lung flotation assay was adapted from previously described methods^34,35^. E18.5 embryos from *minion* ^Δ^*^/^*^+^ *× minion* ^Δ/+^ intercrosses were quickly isolated by cesarean section from humanely sacrificed pregnant females, and were placed on dry Kimwipes. To maintain body temperature, these newborns were incubated by hand and subsequently in a 37 °C chamber. Pups were exposed to normal room air following delivery, and were monitored for at least 1 hour. *minion* ^Δ/Δ^ newborns were uniformly atonic, apneic, and became cyanotic almost immediately after delivery. The majority of *minion*^+/+^ and *minion* ^Δ^*^/^*^+^ mice exhibited normal breathing and demonstrated pink body color indicative of adequate ventilation and perfusion. After at least 1 hour of air breathing, pups were anesthetized, weighed, tailed for genotyping, decapitated, and the lungs were dissected and placed into PBS in 15 ml conical tubes or 2 ml Eppendorf tubes for flotation assay. The lungs were then monitored for more than 15 min, after which they were scored as either floating or sinking. Approximately sixty E18.5 embryos were examined.

### Plasmids and cloning

For the cloning of shRNA constructs, 19-21 nucleotide target sequences were selected using both BLOCK-iT RNAi Designer (Thermo Scientific) and in-house optimized algorithms. For the mouse *minion* mRNA transcripts (GenBank Accession No. NM_001177468.1, NM_001177469.1 and NM_001177470.1), four shRNA target sequences were chosen initially to target all the transcripts, two targeting the coding sequence and two targeting the 3’ UTR. A control sequence was used targeting the firefly (*Photinus pyralis*) *luciferase* gene, which exists in the pGL3 luciferase reporter vector but which lacks similar sequence in the mouse transcriptome. For each shRNA, two 55-59 nt oligonucleotides were designed as shown below and synthesized (IDT). The oligonucleotides, each containing sense and antisense target sequences, a 9 nt intervening hairpin loop, and TTTG at the 5’ends with GATC at 3’ ends for cohesive-end cloning, were annealed. These were then ligated with BbsI/SpeI-digested pGWL-si2/U6 vector. Subsequently using Gateway LR Clonase II Enzyme Mix, these shRNA cassettes were cloned into the vector pLentiLox3.7-GW (pLL3.7-GW), a 3rd generation lentiviral gateway vector that expresses shRNAs under the mouse U6 promoter. A CMV-EGFP reporter cassette was included in the vector to monitor expression.

**Table 2.**
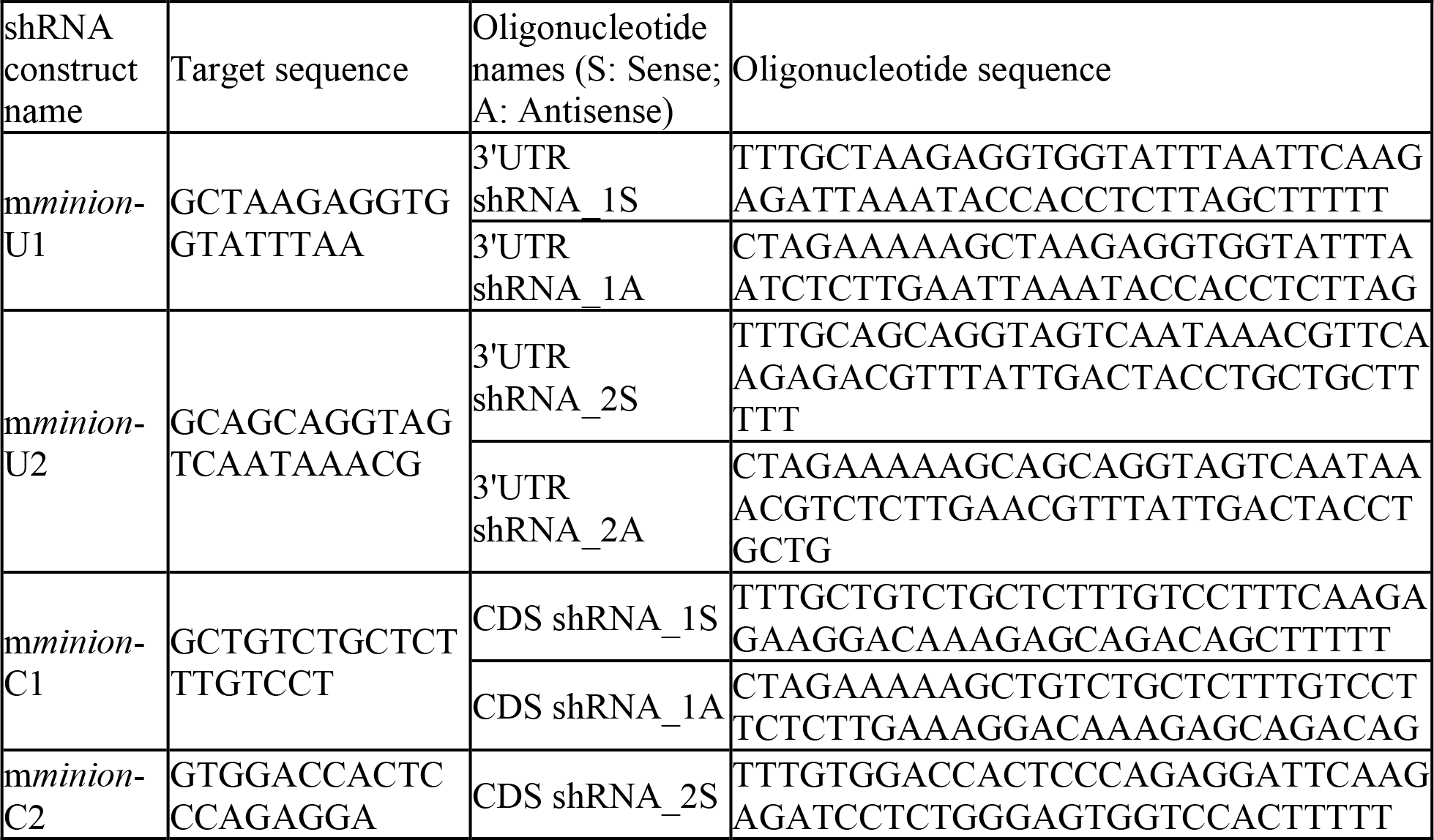
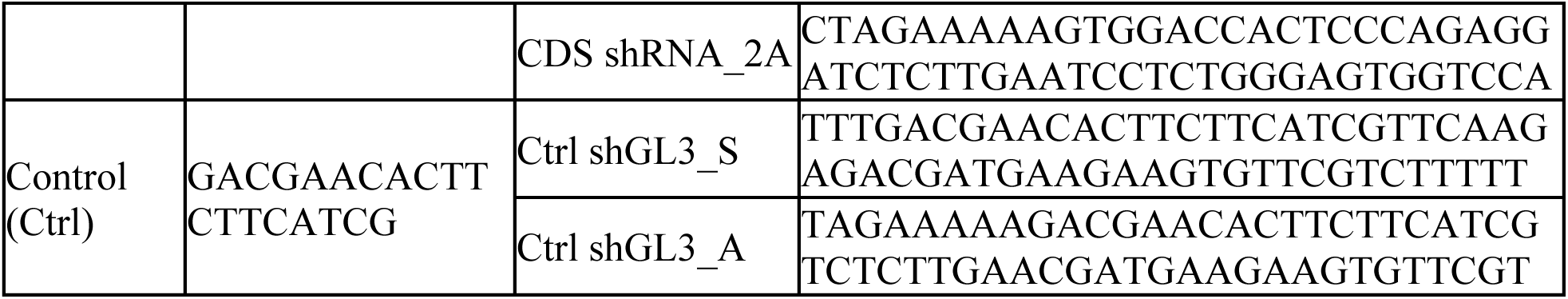
shRNA target sequence and oligonucleotide design information.

For the cloning of cDNA constructs, the coding sequences (CDS) of mouse *minion* (84 aa isoform; CDS and protein accession numbers CCDS50119.1 and NP_001170939.1), human *Minion* ortholog (CDS and protein accession numbers CCDS83093.1 and NP_001302423.1), mouse *tmem8c* (*myomaker;* CDS and protein accession numbers CCDS15823.1 and NP_079652.1), *nanoluc* (control A), *luciferase* (control B), mouse *minion* with 1bp frameshift insertion, human *Minion* with 1bp frameshift deletion, and mouse *minion* truncation (39 aa form) were synthesized (IDT) with attB sites at both ends and a consensus Kozak sequence (5’ GCCACC) before the start codon. For mouse and human *minion* CDS, both untagged and C-terminally 3×FLAG-1×HA-tagged (3F1H) versions were generated. The human *Minion* CDS used here was based on prior genome assembly, whereas the most recent assembly has a C-to-T change and the resulting protein sequence has a R (Arginine) to C (Cysteine) change at the 16^th^ residue. For the *nanoluc* CDS, a C-terminal 1×FLAG tag was added. Using Gateway BP Clonase II Enzyme mix (Thermo Scientific 11789020), the synthesized DNA sequences were cloned into the pDONR221 vector and the sequence-confirmed entry vectors were subsequently cloned into the pCIGAR gateway retroviral vector using Gateway LR Clonase II Enzyme mix. The pCIGAR vector is an MSCV-based bicistronic retroviral vector modified to permit Gateway-mediated insertional recombination of transgenes immediately upstream of IRES-eGFP. In addition, the empty pCIGAR vector (MCS version) was used as another control vector (control C). After expression testing of the two 3F1H-tagged mouse and human *minion* constructs, it was noticed that two bands were detected by Western blot with anti-FLAG antibody (Extended Data Fig.13). This reflected the presence of an extra start codon in-frame with the actual start codon, which was confirmed by sequencing to be inherited from the 3’end of the *CMV* promoter in the pCIGAR vector, giving rise to a product with an additional N-terminal 16 aa, indicated by an asterisk. As a 6-nt consensus Kozak sequence was added before the actual start codon, the intended product is the dominant protein expressed. All other cDNA vectors contained an extra T nucleotide immediately prior to the Kozak sequence, in order to avoid upstream translational initiation. In addition, a pLKO-TREX-On lentiviral MCS vector (kindly provided by Feng Cong, Novartis Institutes for BioMedical Research) was used for making minion expression vector containing C-terminally 3F1H-tagged mouse minion, used for the AP-MS experiments.

### Lentiviral shRNA knockdown assay in primary and transformed myoblasts

Lentiviral particles were produced in HEK 293T cells using a 3^rd^ generation lentiviral packaging system and FuGENE 6 transfection reagent (Promega). Fresh medium was replaced one day after transfection and the supernatant medium was collected on the following day. The medium was briefly centrifuged to remove dead cells, and neat virus was used for QC infection on 293T cells by the reverse infection method with 8 μg/ml polybrene (overnight incubation without spin infection). Analysis of GFP expression by FACS 3 days after infection generally demonstrated a titer of approximately 1 × 10^6^ vp/mL. Neat virus was further concentrated ∼100 fold using a 100kDa centrifugal filter unit (Amicon), aliquoted, and stored at −80°C.

To examine the knockdown efficiency of the mouse *minion* transcript, shRNA-encoding lentiviruses were used to infect C2C12 cells. Viruses were diluted in growth medium containing 8 μg/ml polybrene, and after a brief incubation at 37 °C with C2C12 cells, one round of spin infection was performed at 1100g and 32°C for 1-1.5 hours, using either 24-well or 12-well plates. Based on an estimated viral titer of 1 × 10^8^ vp/ml after concentration (as measured on 293T cells), a virus amount equivalent to MOI30 on 293T cells was used on C2C12 cells. Fresh medium was replaced the next day and GFP^+^ cells were sorted by FACS two days later. Infection efficiency of 70-85% was routinely achieved as judged by FACS analysis, though GFP signal in undifferentiated C2C12 cells was faint by EVOS fluorescence microscopy. Sorted, GFP^+^ C2C12 cells were recovered, expanded and seeded into 6-well plates for *in vitro* differentiation assay. At DM day 4-6, cell lysates were collected and the expression of minion protein was examined by Western blot. The two shRNA constructs targeting the *minion* 3’UTR (U1/U2) were found to reduce minion expression most efficiently (Extended Data Fig.11b), and for subsequent experiments in C2C12 and primary myoblasts, cells were infected with U1 and U2 shRNA viruses only. In order to produce the *minion*^KD^ C2C12 cells, the U1-infected, GFP^+^ sorted cells were reinfected with U2 shRNA virus and resorted for increased GFP signal (top 30-50%). Similarly, wild-type C2C12 cells were infected with the control shGL3 virus (Ctrl) in two rounds and sorted twice to generate the *Ctrl* cells. These *minion*^KD^ and *Ctrl* C2C12 cells were expanded and used for *in vitro* differentiation assay in 384-well and 6-well plates. The *minion*^KD^ cells were also used as the background for *in vitro* reconstitution assay with cDNA-containing retroviruses.

Similar infection steps were followed in order to knock down *minion* in primary myoblasts derived from adult mice, except that 6 μg/ml polybrene was used and fresh medium lacking polybrene was added by the end of the day instead of on the next day, as primary myoblasts appeared more sensitive to polybrene treatment. Due to the difficulty in expanding these infected and sorted primary cells, one round of virus infection was performed using either Ctrl virus or 1:1 ratio mixture of U1 and U2 viruses, and the cells were sorted by GFP signal after 2-3 days. Approximately one-third or one-half the amount of virus used on C2C12 cells was used on primary myoblasts. The sorted primary myoblasts were recovered for 3 days and then seeded into 384-well plates for *in vitro* differentiation assay.

### Retrovirus infection of C2C12 and 10T1/2 cells

The cDNA-encoding retroviruses were made by co-transfecting 293T cells with pCIGAR retroviral vectors and pCL-Eco or pCL-10A1 packaging vectors using FuGENE 6 transfection reagent (Promega). Media was replaced one day after transfection and the supernatant medium was collected on the following day. After a brief centrifugation to eliminate dead cells, the neat virus was used for QC infection in 384-well plate by reverse infection method of NIH-3T3 cells (for pCL-Eco packaged virus) or 293T cells (for pCL-10A1 packaged virus) with 8 μg/ml polybrene (overnight incubation without spin infection). GFP expression was analyzed by FACS 3 days after infection, usually yielding a titer of 0.3-1.2 × 10^6^ vp/ml.

In order to infect C2C12 cells and 10T1/2 cells, neat virus was incubated with 8 μg/ml polybrene at 37 °C for 10 min and then added to the cells, which were then incubated for 15 min at 37 °C. One round of spin infection was performed using 24-well or 12-well plates at 1100 × g and 32 °C for 1-1.5 hours. After 4-6 hours, fresh neat virus with polybrene was added and a second round of spin infection was performed. Fresh medium was added to the infected cells to dilute the polybrene. Media was replaced the following day, and 2-3 days later, the infected cells usually exhibited very strong GFP fluorescence by EVOS microscopy. Infection efficiency using this two-round infection method was generally >95% over a large range of viral concentrations, therefore FACS was not generally necessary. The resulting GFP signal was used to mark the boundary of cells, in addition to phase contrast imaging and nuclear markers (Extended Data Figs. 18, 20); moreover, in cells that were fixed with 4% PFA but not permeabilized with detergents, the GFP signal was higher in the nuclear region, allowing unambiguous delineation of both the cell boundary and nuclei.

For each round of infection, a viral amount equivalent to MOI 3-6 on 3T3 or 293T cells was used for *minion*^KD^ C2C12 cells, and the infected cells were expanded and used for *in vitro* reconstitution assay in differentiation medium. For co-expression and cell-mixing experiments on wild type 10T1/2 cells and C2C12 cells, a viral amount equivalent to MOI 2-4 on 3T3 or 293T cells was used. For each comparison, the viruses made with the same packaging plasmid and at similar MOI were used for infection. The experiments were repeated with different types of control viruses (Luciferase, NanoLuc-FLAG, empty vector). For co-expression and mixing experiments, cells were infected with one type of cDNA retrovirus first using the 2-round spin infection protocol described above, expanded for several days, then reinfected with a second cDNA-encoding retrovirus using the same method. On the day following the final infection, the cells were labeled with different dyes, mixed, and seeded into 384-well or 24-well plates as described below.

### Affinity purification and mass spectrometry (AP-MS) analysis

C2C12 myoblasts expressing both C-terminally 3×FLAG-1×HA-tagged mouse minion and empty control were seeded in growth medium, then incubated in differentiation medium for 3.5 days with 2 μg/ml Doxycycline to induce the expression of exogenous minion protein from pLKO TREX-ON lentiviral cDNA vector. These cells were then used in one-step anti-FLAG immunoprecipitation. Cytoplasmic and membrane fractions were prepared using the ProteoExtract Native Membrane Protein Extraction Kit (Millipore 444810). Cytosolic and membrane fractions were subjected to affinity purification using anti-FLAG M2 affinity gel (Sigma F2426) with 1 hour incubation, and the immunopurified protein complexes were eluted by the addition of FLAG peptide (Sigma F4799). The eluate was precipitated by trichloroacetic acid and washed twice with acetone. The precipitates were dissolved in digestion buffer (8M Urea in 100 mM Tris pH 8.5), reduced, alkylated and digested as described previously^36^. The digested peptide mixture was desalted using C18 pipette tips (Thermo Scientific) and fractionated online using a 75 μM inner diameter fused silica capillary column, with a 5 μM pulled electrospray tip and packed in-house with 16 cm of C18 reversed phase spherical silica particles of 3.0 μM.

An Ultimate 3000 nano LC system (Thermo Scientific) coupled to a Q Exactive mass spectrometer (Thermo Scientific) was used to deliver the linear acetonitrile gradient using buffer A (0.1% formic acid water) and buffer B (0.1% formic acid water, 100% ACN) starting from 5% buffer B to 35% over 70 min at a flow rate of 200 nL/min, followed by a 5 minute ramping to 95% acetonitrile and a 5 minute hold at 95% buffer B. The column was re-equilibrated with 2% buffer B for 2 minutes before the subsequent run. Each sample was analyzed in triplicate technical replicates on LC-MS/MS.

MS/MS spectra were collected on a Q-Exactive mass spectrometer (Thermo Scientific) as described previously^37,38^. All MS/MS raw spectra files were processed using Proteome Discoverer 1.4 (Thermo Scientific), and the spectra from each raw file were converted to MASCOT generic files (MGF) for dataset searches using licensed Mascot Daemon (Matrix Science, London, UK; version 2.4.1). The MS/MS data was searched against the Uniprot database downloaded on April 1^st^, 2015 (selected for *Mus musculus* with 16715 entries) including a decoy database. The carbamidomethylation of cysteines was set as a static modification with a mass of +57.02156 Da. The specificity of digestion was set for trypsin allowing three missed cleavages. The mass tolerances in MS and MS/MS mode were set to 15 ppm and 0.8 Da, respectively. Peptide and protein identifications were validated using Scaffold version 4.6.2 (Proteome Software Inc., Portland, OR). Peptide and protein identifications were accepted if they were detected with ≥95.0% and ≥99.0% probabilities respectively by the Scaffold local false discovery rate algorithm, and required at least 2 identifiable unique peptides^39^. Fisher’s exact test p-value in Scaffold were calculated according to a model previously described^40^. The protein classes of significantly enriched proteins were identified and visualized using the Panther classification system (http://www.pantherdb.org)^32^.

### Cell labeling and mixing experiments

The performance of a series of cell-permeant fluorescent dyes was tested on 10T1/2 fibroblasts and C2C12 myoblasts over multiple dilutions and using different labeling methods, with signal strength and pattern monitored continuously by microscopy for at least four days. CellTracker Deep Red dye (Thermo Scientific C34565, 1:250 final dilution) and CellTrace Violet dye (Thermo Scientific C34571, 1:500 final dilution) were selected for subsequent experiments, and were observed to label the cytoplasmic and nuclear regions of mononuclear cells. However, in fused multinuclear cells (Fig. 4f, Extended Data Fig. 20), the CellTrace Violet dye exhibited a strong enrichment in nuclei which had been originally labeled with the dye, and did not diffuse into other non-violet-labeled nuclei, while the CellTracker Deep Red dye demonstrated perinuclear enrichment and was helpful in recognizing the cell boundary. These features allowed facile quantification of fusion efficiency (Fig. 4g).

Cell labeling was performed according to manufacturer’s instructions with slight modifications. Cells were trypsinized, centrifuged, washed once with PBS, transferred to a 2 ml Eppendorf tube, centrifuged, and resuspended in PBS at a concentration of 0.8-1 × 10^6^ cells/ml. 2x cell labeling solution was prepared separately in PBS with fluorescent dyes and was mixed well by brief vortexing. Equal volumes of the 2x labeling solution and the cell suspension (usually 250 μL each) were mixed and incubated at 37 °C for 40-45 min with occasional mixing. The labeled cells were then mixed with 1 ml fresh medium, incubated at 37 °C for 5 min and centrifuged at 150 × g. At this point, 3 more washes of the labeled cells were performed using fresh medium, incubation at 37 °C for 10-15 min, and centrifugation at 150 × g. After the last wash, the cells were counted and diluted to the concentration needed for the final mixing experiment in a 384-well or 24-well plate, and were incubated at 37 °C for 30 min. These additional washes and incubation steps were used to eliminate remaining unbound dye in the cell suspension and on the cell surface, which was critical for cell mixing experiments performed on the same day. After incubation, cells labeled with Deep Red dye were mixed with those labeled with Violet dye at a 1:1 ratio, and were seeded into 384-well or 24-well plates, with a range of concentrations tested from 800-4000 cells/well. Cells in 384-well plates were fixed at different time points (24-48 hours) with 4% PFA for 10 min at room temperature, and were washed with PBS before imaging. Growth medium containing 10%-15% FBS was used for all experiments in both 10T1/2 and C2C12 cells.

### Differentiation and fusion indices

Several indices were used to examine the fusion efficiency in C2C12 cells as well as adult and embryonic primary myoblasts during *in vitro* differentiation. The differentiation index shown in C2C12 (Fig. 3e) and adult primary myoblasts (Fig. S12b) was calculated as the fraction of nuclei contained within all MHC^+^ cells, including both mononuclear and multinuclear cells, as compared with the number of total nuclei within each 20x image acquired by IXM confocal high-content imaging. At least 6 separate fields from independent replicate wells were quantified for each genotype each experimental repeat. The fusion index shown in C2C12 (Fig. 3f) and adult primary myoblasts (Extended Data Fig. 12c) was calculated as the fraction of nuclei contained within MHC^+^ myotubes which had three or more nuclei, as compared to the number of total nuclei within each 20x image taken by IXM imaging. At least 6 separate fields from independent replicate wells were quantified for each genotype. The fusion index shown for embryonic primary myoblasts (Fig. 3b) was calculated as the fraction of nuclei contained within Desmin^+^ myotubes having three or more nuclei, as compared to the number of nuclei within all Desmin^+^ cells on each 20x IXM image. At least 6 separate fields from independent replicate wells were quantified for each genotype each experimental repeat. Since fibroblasts still existed in these early-passage primary cultures, only the total nuclei in Desmin^+^ cells were included for quantification. Another index used to examine the fusion efficiency in C2C12 cells (Fig. 3g) and adult primary myoblasts (Extended Data Fig. 12d) was focused on myotubes. Myotubes were binned into 3 to 4 subgroups based on the number of nuclei contained within each tube: 2 nuclei, 3-5 nuclei, 6-10 nuclei, or more than 10 nuclei (Fig. 3g). The fraction of each subgroup was calculated for each genotype in comparison to total myotube number. At least 6 separate fields from independent replicate wells were quantified for each genotype. Please find more details about the numbers in the figure legends.

For co-expression experiments in 10T1/2 cells, two indices were used to examine fusion efficiency. The first fusion index (Fig. 4e) was based on the experiment shown in Extended Data Fig. 18, and was calculated as the percentage of nuclei within GFP^+^ syncytia containing 3 or more nuclei, as compared with the number of total nuclei. At least four separate 10x IXM images from different replicate wells were used for quantification of each genotype. In order to examine the ability of minion-only or myomaker-only expressing cells to fuse with minion and myomaker co-expressing cells as well as to further confirm the formation of multinuclear syncytia by fusion instead of incomplete cytokinesis, a second index was used (Fig. 4g) to calculate the fusion efficiency between Deep Red dye-labeled cells and Violet dye-labeled cells in three different combinations (Fig. 4f, Extended Data Fig. 20). In these combinations, all of the Deep Red dye-labeled cells had co-expression of myomaker and minion, no matter which protein was expressed first by infection, and these cells were able to fuse to themselves, becoming Deep Red^+^ syncytia. The Violet dye-labeled cells expressed 1) myomaker and minion (either myomaker or minion was expressed first); 2) myomaker only (together with empty vector control); 3) minion only (together with Luciferase control). The fraction of Deep Red^+^ syncytia (containing 3 or more nuclei) containing one or more Violet dye-labeled nucleus was compared with the total number of Deep Red^+^ syncytia on each 10x IXM image. At least 6-12 separate fields from independent replicate wells for each experimental repeat were quantified for each combination.

### Statistical analysis

Sample size was not predetermined statistically and no specific blinding method or randomization was applied. Each value reported represents the mean±s.d. of more than two independent replicates as described for each experiment. Replicate types utilized include: independent experimental (biological) replicates; individually treated cells with viruses or compounds; mice from the same or different litters; and tests or assays run on the same sample multiple times (technical replicates). Quantitative data were analyzed by unpaired two-tailed Student’s *t*-test with Welch’s correction and without assumption of equal standard deviations. All statistical analyses were performed using GraphPad Prism software. *P* < 0.05 was considered statistically significant. *P* < 0.05 was marked with an asterisk (*) and *P* < 0.001 was marked with double asterisks (**).

### Data availability

Source data for proteomic studies is provided in Extended Data Table 1. Raw proteomic data are also available on request.

